# Control of walking direction by descending and dopaminergic neurons in *Drosophila*

**DOI:** 10.1101/2025.07.22.666129

**Authors:** Stefan Dahlhoff, Sander Liessem, Fathima Mukthar Iqbal, Adrián Palacios-Muñoz, Federico Cascino-Milani, Mert Erginkaya, Aleyna M. Diniz, E. Axel Gorostiza, Ansgar Büschges, Jan Clemens, Jan M. Ache

## Abstract

Animals need to fine-control the speed and direction of locomotion to navigate complex and dynamic environments. To achieve this, they integrate multimodal sensory inputs with their internal drive to constantly adjust their motor output. This integration involves the interplay of neuronal populations across different hierarchical levels along the sensorimotor axis – from sensory, central, and modulatory neurons in the brain to descending neurons and motor networks in the nerve cord. Here, we characterize two populations of neurons that control distinct aspects of walking on different hierarchical levels in *Drosophila*. First, we use *in-vivo* electrophysiological recordings to demonstrate that moonwalker descending neurons (MDN) integrate antennal touch to drive changes in walking direction from forward to backward. Second, we establish DopaMeander as an important component in the control of forward walking through a combination of optogenetic activation, silencing, connectomics, and *in-vivo* recordings. These dopaminergic modulatory neurons drive forward walking with increased turning, and the activity of individual neurons is correlated with ipsiversive turning. Hence, MDN and DopaMeander control opposite regimes of walking on different hierarchical levels. Computational models reveal that their activity predicts key parameters of spontaneous walking. Moreover, we find that both MDN and DopaMeander are gated out during flight. This suggests that neuronal populations across levels of control are modulated by the behavioral state to minimize cross-talk between motor programs.

## Introduction

Although it appears effortless, controlling the speed and direction of walking is a complex motor task that requires coordinated activity across muscles, joints, limbs, and body segments. This adaptability is essential for survival, supporting behaviors such as foraging, courtship, and predator evasion^1–6^. In both insects and vertebrates, motor control networks are distributed across the brain – which integrates sensory input, internal states, and higher-level decisions – and the ventral nerve cord (VNC, in insects) or spinal cord (in vertebrates), which execute motor commands^2^. In *Drosophila*, motor commands are transmitted from the brain to the VNC by a small population of approximately 1,300 descending neurons (DNs)^7^, which link 140,000 brain neurons to 15,000 VNC neurons^8–11^.

The neuronal control of walking is distributed not only across the brain and VNC but also across hierarchical levels within the brain. For example, feature-detecting neurons in the optic lobe^12–14^ and antennal mechanosensory neurons distribute their output across central and descending pathways^15^ to influence walking behavior^16–19^. At a higher level, modulatory neurons — including octopaminergic, dopaminergic, and different peptidergic systems — adjust brain circuit activity to increase or decrease walking speed and motivation^2,5,20^. Thus, classical sensorimotor pathways link sensory networks in the brain to motor networks in the VNC, while an upstream modulatory layer tunes the internal drive to walk. This circuit architecture is similar in insects and vertebrates. Here, we characterize two neuronal populations that control distinct aspects of walking and operate on different levels within the motor hierarchy: a population of DNs that directly connect sensory circuits in the brain to motor circuits in the VNC, and a population of upstream modulatory neurons.

The overall organization of descending pathways is well-described in *Drosophila* and other insects^2,10,21^, and several identified DNs have been shown to regulate specific walking features^22^. These include DNs involved in initiating^14^ and terminating^23^ walking, as well as those that control direction^24–26^ and speed^14^. Some DNs have been labeled ‘command-like neurons’ because they are both necessary and sufficient for specific behavioral outputs. However, adaptive walking requires coordinated activity across the DN population^2,26^. To understand this population-level coding, we must quantify DN activity across behavioral contexts. While many recent studies have used genetic activation or silencing to infer DN function, far less is known about the dynamic activity changes of DN populations in behaving animals. Here, we provide a detailed *in-vivo* characterization of a command-like DN population in *Drosophila* that controls the switch from forward to backward walking – the moonwalker descending neurons (MDNs)^13,27,28^.

The MDN population comprises four neurons, two per hemisphere. MDNs reliably induce sustained backward walking when artificially activated, and they are necessary for backward walking in confined environments^27^. Their downstream VNC targets act during both stance and swing phases of the hind leg to reverse the forward-walking motor pattern and generate backward walking^28^. Unilateral MDN activation results in contraversive backward turns, and the number of active MDNs positively correlates with the distance walked backward^13,29^. Coactivation of MDNs with moonwalker ascending neurons (MANs), which are thought to inhibit forward walking, enhances the consistency of backward walking ^27^. Notably, MDNs also mediate backward locomotion in *Drosophila* larvae^30,31^, suggesting a conserved motor function for MDN across distinct body plans and modes of locomotion.

MDNs integrate multimodal sensory input to guide directional changes: they receive input from LC16 visual projection neurons^13^, which are responsive to visual looming stimuli^12,32^, and from touch-sensitive neurons in the front legs^29^. These inputs trigger backward walking away from the stimulus, presumably aiding in predator evasion and obstacle avoidance. Population-level imaging experiments show that MDN activity correlates with backward walking^33^. Despite their central role in motor control – and their status as one of the best-characterized DN types – the precise *in-vivo* activity dynamics of MDNs during behavior are unknown. Therefore, it remains unclear under which naturalistic conditions MDNs get activated, and how MDN activity translates into behavior. Here, we show that MDNs integrate antennal mechanosensory input to drive backward walking and quantify how MDN activity relates to spontaneous changes in walking direction. This reveals antennal touch as a previously unrecognized sensory context for backward locomotion in *Drosophila*, which is shared with other insects^16,34^.

While *Drosophila*, like most animals, employ forward walking as their primary mode of locomotion, its neuronal underpinnings remain poorly understood, with only a few central and descending neurons known to contribute^14,23–25^. To address this gap, we investigated a genetic driver line (SS02553), which was previously shown to drive strong increases in forward walking speed as part of an extensive optogenetic activation screen^22^, making it an interesting contrast to MDN. Here, we combined genetic activation and silencing approaches with electrophysiological recordings and connectome analysis to demonstrate that this forward walking phenotype is driven by a unique type of dopaminergic modulatory neuron in the central brain, DopaMeander, which drives forward walking and ipsiversive turning.

Leveraging the fact that we have identified neurons which control walking on different hierarchical levels – central modulatory and descending neurons – we asked whether neuronal populations that control walking also contribute to flight. We hypothesized that involvement in multiple behaviors might depend on a neuron’s position within the motor control hierarchy – with modulatory neurons acting as generalists and DNs being more behavior-specific. To test this, we examined MDN and DopaMeander activity during flight, probing their functional roles across behavioral states. This allows us to explore how walking is modulated by different neuronal populations and whether these circuits are repurposed or bypassed during other behaviors.

## Results

### MDN drives antennal touch-mediated backward walking

In numerous insect species, mechanical stimulation of the antennae induces backward walking^16,34^, providing a relevant ethological context for switches in walking direction. To test if *Drosophila* exhibits a similar response, we stimulated flies tethered on a spherical treadmill with gentle, frontal touches to the antennae, which resulted in posterior deflection, and recorded their walking trajectories (Figure 1A). Antennal stimulation resulted in brief episodes of backward walking, reaching maximum speed 300 ms after the onset of stimulation. Simultaneously, the angular velocity increased towards the contralateral side relative to the stimulated antenna (Figure 1B). This indicates that flies executed a backward turn away from the stimulated antenna (Figure 1C).

**Figure 1:**
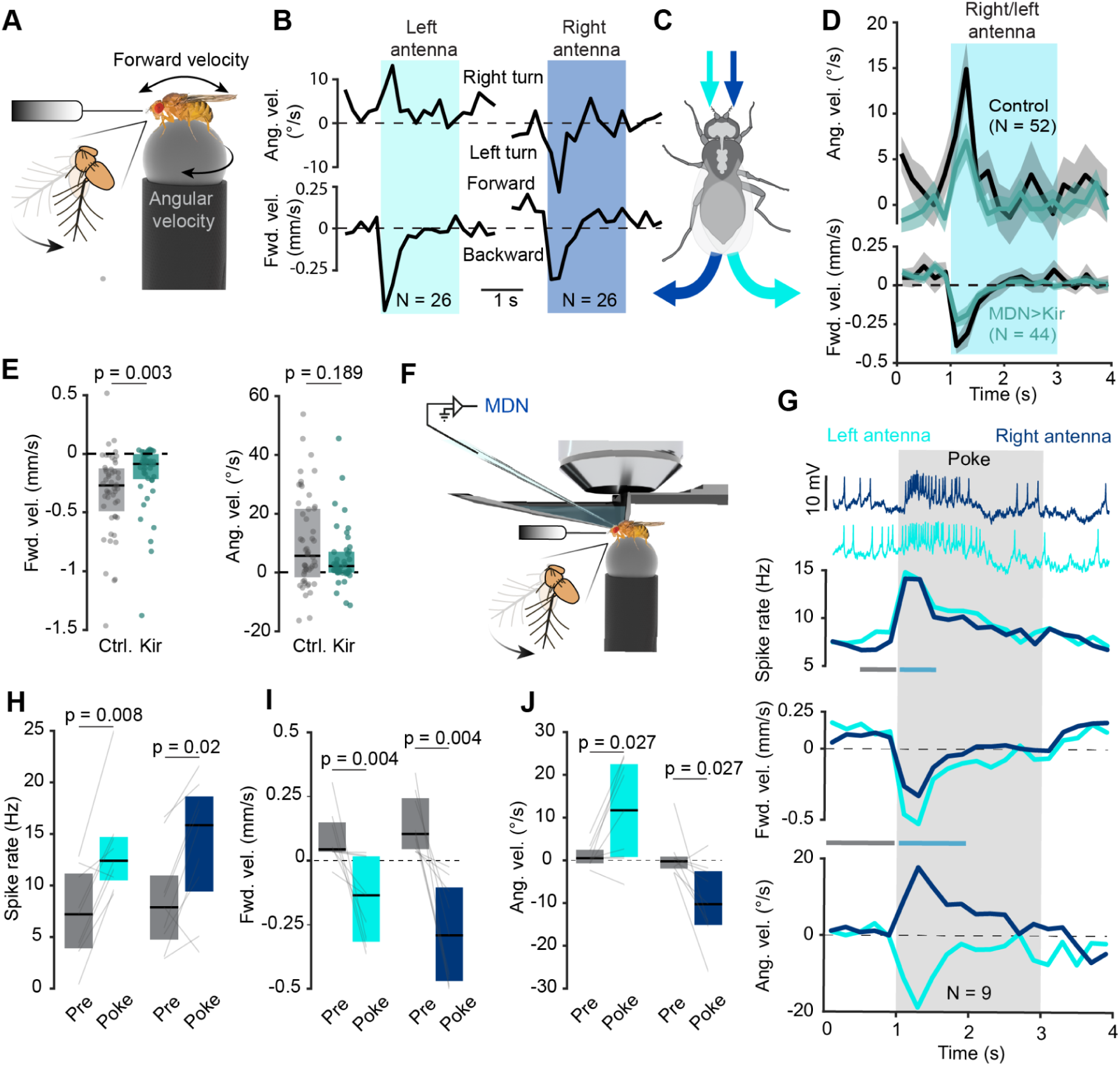
MDN integrates bilateral antennal mechanosensory input to drive backward walking. A) Experimental setup for antennal mechanical stimulation in tethered walking flies. B) Angular (top) and forward velocity (bottom) during antennal stimulation (colored rectangles) in 200 ms bins. Data averaged across 260 trials per antenna in 26 flies. C) Antennal touch elicits backward walking directed away from the side of the stimulation. D) MDN silencing reduces antennal touch-driven backward walking. Angular and forward velocity during antennal stimulation (colored rectangle) pooled over the left and right antenna for control (MDN>GFP, black, N=52) and silenced MDN>Kir2.1 flies (teal, N=44). Angular velocity from the right antenna was mirrored before averaging. Shading indicates standard error. E) Quantification of pooled forward and angular velocity during the first 500 ms of antennal stimulation for MDN>GFP (ctrl.) and MDN-silenced flies (Kir2.1). Points represent mean velocity for each fly, boxplots show median ± IQR. One outlier in control at 1.48 mm/s out of scale for clarity of inspection. P-values were calculated using the Wilcoxon signed-rank test. F) Experimental setup for electrophysiological recording during single antennal touch stimulation. G) MDN activity and walking trajectories during antennal touch stimulation (grey rectangle). From top to bottom: MDN recording during stimulation of right (dark blue) and left (light blue) antenna. Traces, from top to bottom, show the mean spike rate, the mean forward velocity, and the mean angular velocity averaged across ten trials for each antenna from all flies (N=9). Data was binned in 200 ms windows. H) Quantification of MDN activity during antennal touch. Spike rate was analyzed 500 ms before (pre) and in the first 500 ms of stimulus (poke). Data shown as median ± IQR. I) Quantification of forward velocity during MDN recordings in response to antennal stimulation. Forward velocity was averaged during 1 s before (pre) and in the first 1s of the stimulus (poke). Other details as in H. J) Angular velocity changes during MDN recording in response to antennal touch. Other details as in I. P-values in (H-J) were calculated using the Wilcoxon rank-sum test.

Since MDN is firmly established to control backward walking in *Drosophila* and capable of driving backward turning upon unilateral activation^13^, we explored the involvement of MDNs in antennal touch-mediated backward turning. For this purpose, we repeated the antennal stimulation experiments in flies in which the MDN population was silenced by expressing Kir 2.1 under an MDN-specific driver line (MDN-1^27^). Data from all antennal stimulation trials across the left and right antenna was pooled by mirroring angular velocities during stimulation of the right antenna so that right and left turns had the same sign (Figure 1D). The backward walking velocity during antennal stimulation was significantly reduced by 66% in MDN-silenced flies compared to controls (−0.27 mm/s vs. −0.09 mm/s, p = 0.003). This was accompanied by a reduction in angular velocity, which, however, was not significant due to the large variability in turning velocities in control flies (0.31 mm/s vs. 0.11 mm/s, p = 0.19) (Figure 1E). Hence, MDN is required for full expression of antenna-mediated backward walking. Interestingly, MDN silencing affected primarily the backward walking component and had weaker effects on the turning component. In summary, MDNs are required for the backward turning response to antennal touch. The fact that flies with silenced MDNs still walked backwards implies that there could be a parallel pathway driving backward walking independent of MDN. Alternatively, MDN silencing via Kir 2.1 might have been incomplete.

### MDN integrates antennal mechanosensory input

While the role of MDN in controlling backward walking has been firmly established through behavioural experiments, MDN has not been characterized extensively in behaving animals. To close this gap, we performed *in-vivo* patch-clamp recordings in behaving flies to understand how MDN activity drives changes in walking direction. First, we investigated how MDN controls backward walking in response to antennal touch (Figure 1F).

MDNs exhibited a baseline spike rate of 8.5 Hz with a median spike amplitude of 6.5 mV when flies were tethered, and the legs were in contact with the treadmill. The resting membrane potential was −39 mV (without subtraction of a 13 mV junction potential^35^). Mechanical stimulation of the antenna resulted in a sharp increase in MDN spike rate within 100 ms, which matched the temporal dynamics of the backward walking response so that the maximal activity of MDN occurred during the backward walking bout (Figure 1G). The MDN spike rate remained elevated for about 500 ms during antennal stimulation. After antennal stimulation, MDN activity and walking behavior returned to baseline (Figure S 1A-C).

Importantly, each MDN responded to stimulation of the left and the right antenna, so that a given MDN was active during backward turning towards the left and the right side, and hence ipsi- or contraversive with respect to its soma location (Figure 1G). This observation held across individuals – each MDN was roughly equally responsive to stimulation of the right and the left antenna (Figure 1H), with stimulation of the left antenna driving a change in MDN spike rate from 7.2 Hz to 12.4 Hz, and the right from 7.8 Hz to 15.8 Hz, on average. The increases in MDN spike rate occurred during bouts of backward walking (Figure 1I), directed either towards the left or right direction away from the stimulated antenna (Figure 1J), consistent with our earlier behavioral experiments. This strongly suggests that individual MDNs do not contribute to the direction of backward turning during antennal stimulation. To test this, we quantified if individual MDNs showed a preference for stimulation of the right or left antenna. A contrast analysis comparing the response to stimulation of the left and right antenna for each MDN revealed a very weak preference, so that each MDN encodes little information about which antenna was stimulated (Figure S1D). In contrast, the same analysis for the angular velocity of backward turning revealed a much stronger preference for right turns in response to left antennal touch and left turns in response to right antennal touch (Figure S1E). Hence, the behavioural response was lateralized, but the MDN population activity was not.

Since individual MDNs responded equally strongly to stimulation of the left and right antenna, MDN cannot drive the turning component of the behavioral response to antennal stimulation – the MDN spike rate is positively correlated with turning in both directions. Together with the observation that MDN silencing does not significantly affect the turning component of antenna-mediated backward walking, this clearly shows that the MDN population drives straight backward walking, and not turning, in the context of antennal mechanosensory stimulation. Elegant behavioural experiments by Sen et al.^13^ established that optogenetic activation of individual MDNs drives ipsiversive backward turning. Hence MDNs can drive backward turning, but our electrophysiological recordings in behaving flies show that, in the context of antennal stimulation, they do not. Since flies turned away from the touched antenna nonetheless, we postulate that a second, MDN-independent pathway must drive the turning component in this context.

To test whether individual MDNs also receive bilateral input from the visual system, we performed a series of visual stimulation experiments while recording MDNs in stationary flies. Since it is not possible to distinguish between left and right MDNs based on soma location during patch-clamp experiments, we stimulated both eyes and recorded from several MDNs subsequently in individual flies. In general, we observed robust responses with increasing spike rates as soon as visual stimuli appeared on the LED arena surrounding the fly (Figure S1F-H). These responses persisted after antennal ablation, suggesting they are not driven by antennal movements which might be triggered by visual stimulation (Figure S1H). Visual responses to widefield motion and moving objects were weak but relatively consistent under these conditions (Figure S1I-N). For instance, regressive visual motion, which occurs during backward walking, only had weak effects on MDN activity (Figure S1J). The strongest visual responses were elicited by frontal approaching bars (Figure S1M-N), in line with the sensitivity of MDNs to approaching or looming objects^13^. Importantly, recording from all four MDNs subsequently in a single fly revealed that individual MDNs responded to visual stimulation in both hemispheres with no differential tuning in terms of rotational direction and other parameters of visual stimulation (Figure S1O). This suggests that all four MDNs were tuned to the same directions of motion, and therefore not to visual motion cues which are experienced during and drive turning. We conclude that the MDN population integrates bilateral information from the visual and antennal mechanosensory system to control backward walking.

### A neuronal substrate for forward and curve walking

MDN serves as a command-like neuron that initiates and controls backward walking. To gain a more comprehensive understanding of the neuronal control of walking, we screened genetic driver lines implicated in controlling forward walking, to identify neurons that operate in the opposite control regime of MDN. To this end, we analyzed behavioral effects of DN lines implicated in forward walking by a previous, extensive optogenetic DN activation screen^22^. We selected the most-promising DN candidates that were linked to increases in walking speed by Cande et al. and featured a posterior soma location (DNp)^10^, so that they were accessible for *in-vivo* patch-clamp recordings. As expected, activation of these DNp lines also increased the walking speed in our behavioral assays.

The split-Gal4 driver line SS02553, which labels DNp17 (among other neurons), had the most prominent and robust impact on forward walking. We therefore focused on this driver line in subsequent experiments. DNp17 forms a population of 6 neurons per hemisphere, which arborize ipsilaterally in the brain and descend ipsilaterally into the VNC^10^, where they primarily innervate wing and haltere related neuropils. First, we confirmed the phenotype associated with SS02553 via optogenetic activation in our Universal Fly Observatory (UFO) setup (Figure 2A). Activation drove consistent increases in forward walking speed, leading to longer walking trajectories during the period of red light exposure compared to ‘empty’ controls^10^ with no expression in the nervous system (Figure 2B, Video S1). SS02553 activated flies increased their walking speed immediately after the onset of activation, and kept walking faster than controls throughout the activation period (Figure 2C). This became even more evident when averaging over all flies and across activation light intensities (ranging from 0.4 to 4 mW/cm^2^), revealing significantly faster forward walking compared to controls exposed to the same stimulus protocol (Figure 2D).

**Figure 2:**
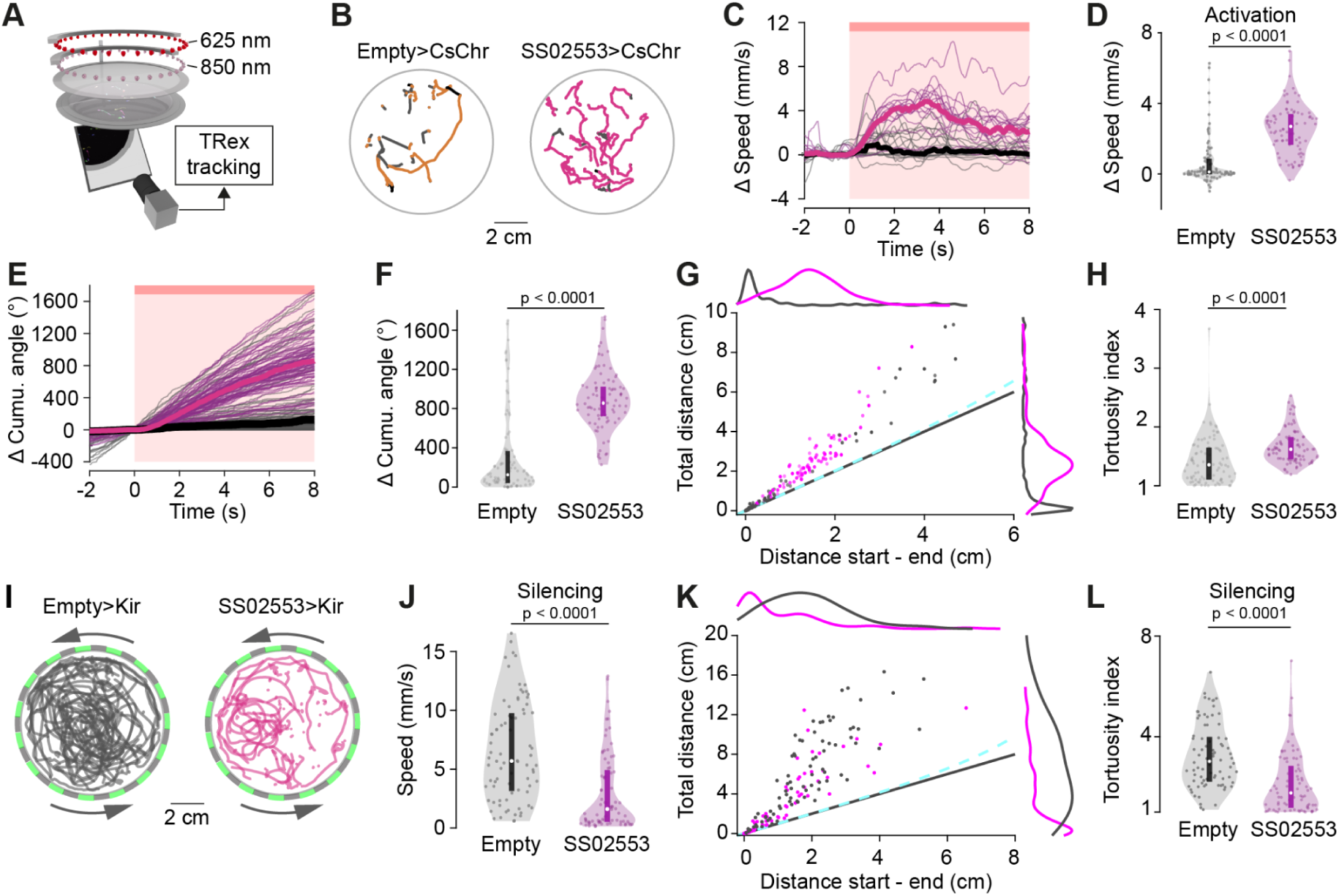
SS02553 drives forward walking with increased turning. A) Schematic of UFO setup for optogenetic activation. B) Left, example trajectories from empty>CsChrimson controls (N=18) and SS02553>CsChrimson flies (N=17) before (black), during (orange/magenta) and after (grey) a single activation pulse. C) Walking speed during 9 activation trials for controls (grey, N=18) and SS02553>CsChrimson (magenta, N=17). Median baseline speed during 2 s before activation was subtracted. Light intensity, 2.3 mW/cm^2^. Thin lines, averages for each fly across 9 trials, thick lines, median across all flies. D) Walking speed of empty>CsChrimson (grey, N=18 flies, 5 intensities each) and SS02553>CsChrimson (magenta, N=17 flies, 5 intensities each) averaged over 9 activation trials per intensity. Time window as in C. E) Cumulative walking angle of controls (grey, N=18 flies, 5 intensities each) and SS02553>CsChrimson (magenta, N=17, 5 intensities each) averaged over 9 activation trials per intensity. Each trace was baseline subtracted at 0 s. Thick lines, median across all flies. F) Each dot represents the final value at the end of the 8 s window in (E), shown with the distributions. G) Comparison of total distance walked and distance between the start and end point during the first 8s of activation for control and SS02553>CsChrimson flies. Each dot represents the mean across all 9 activation trials for one fly at one light intensity. Opacity of dots increases with activation light intensities (0.4, 1.4, 2.3, 3.2, 4.1 mW/cm^2^). Distributions show the normalized kernel density for total distance walked and distance between start and end point. Dashed line indicates theoretical function of distance walked if a fly followed the arena wall, which cannot explain increased turning rate. Identity line shown in black. H) Tortuosity index, calculated by (total distance walked/distance start to end). A higher tortuosity index indicates more turning during walking. I) Example trajectories of freely walking control (gray) and SS02553>Kir2.1 silenced flies (magenta) under optomotor stimulation. In these experiments, a grid of 15 mm wide vertical LED stripes moved counterclockwise at 15 cm/s around a group of 4 starved male or female flies to stimulate walking and turning. The experiment was repeated 20 times, so that N=80 flies per genotype. J) Mean forward walking speed of control and SS02553>Kir2.1 flies. Each individual dot represents the mean forward walking speed for each fly, N=80 flies per condition. K) SS02553 silencing reduced forward walking and turning compared to empty controls. Plot details as in G. L) The tortuosity index of SS02553 silenced flies was significantly smaller than of empty>Kir2.1 controls. P-values in all panels are calculated via Wilcoxon rank sum test.

Interestingly, optogenetic activation of SS02553 also increased curve walking: flies did not simply walk faster than controls, but they also walked along a meandering trajectory (Figure 2B). First, we quantified this by calculating the cumulative turning angle of all flies averaged across all activation trials (Figure 2E). While the cumulative angle was flat before activation, SS02553 activation led to a steady increase of the cumulative angle, indicating the flies consistently turned during forward walking. Control flies, in contrast, did not exhibit an increased cumulative angle over time (Figure 2F). However, it is somewhat expected that flies which walk more also turn more, simply because they show more activity.

To quantify this effect while accounting for systematic differences in walking speed between SS02553 activated flies and controls, we approximated the tortuosity of each walking trajectory by quantifying the total path length relative to the distance between the start and end point during optogenetic activation (Figure 2G). This analysis revealed that SS02553 activation led to increased walking distance and increased turning. The increase in turning resulted in significantly increased tortuosity indices (total distance/distance start-end, Figure 2H), verifying that SS02553 activation leads to significantly increased turning during walking. The light intensity, and hence the spike rate of activated neurons, only had a small effect on the amount of turning during activation, and a negligible effect on the walking speed (Figure S2A & B). Overall, activation of SS02553 elicits forward walking with increased curve walking – in contrast to MDN, which drives straight backward walking. Notably, the increased turning, or meandering, occurred even though we activated the neurons labeled by the SS02553 line bilaterally, suggesting a circuit mechanism that generates temporal asymmetries in the activity patterns on the right and left side of the CNS.

Next, we asked if the neurons labeled by SS02553 are necessary for fast forward walking and turning. To test this, we used a UFO setup surrounded by an LED arena for visual stimulation (Figure 2I). We presented a grid of vertical stripes moving counterclockwise around the flies, which drives an optomotor response that reinforces walking on circular trajectories (Figure 2I)^36^. Control flies exposed to this stimulus showed increased walking activity with high speeds, and often walked along circular trajectories in the center of the arena (Figure 2I, J). Walking activity and walking speed were significantly reduced in SS02553-silenced flies (Figure 2I, J, Figure S2C), which also exhibited fewer circular trajectories (Figure 2I). Accordingly, their cumulative angle during walking was strongly and significantly reduced compared to controls (See Figure S2D). Generally, control flies walked and turned much more when they were starved^37^ and presented with an optomotor stimulus, as expected (compare Figure 2K and 2G). However, SS02553-silenced flies were consistently shifted towards the lower right in the behavioral space defined by the total walking distance vs. the start-to-end distance, suggesting that they walked less and made fewer turns when walking (Figure 2K). Accordingly, the tortuosity index of SS02553-silenced flies was significantly lower than in controls, confirming that SS02553-silenced flies turned less (Figure 2L). Overall, our activation and silencing experiments showed that neurons in the SS02553 line drive forward walking and turning, and are required for flies to fully express the optomotor response during free walking. Hence, the neurons labelled by the SS02553 line – presumably DNp17 – and MDN populations control complementary parameters of forward and backward walking and turning.

### DopaMeander drives the walking phenotype of SS02553

The behavioral experiments revealed that the SS02553 driver line clearly modulates walking, in that neurons labelled by this line increase forward speed and turning. While most of the DN split-Gal4 driver lines published are very specific and label primarily one population of DNs each, this is not the case for SS02553, which labels several other neuron types in addition to DNp17^10^. To test if the activation and silencing effects on walking we observed were driven by the DNp17 population, we selectively recorded from DNp17 via patch-clamp, in tethered walking flies, using the same approaches as for MDN recordings. DNp17 was spontaneously active and fired action potentials at 17 Hz at rest. However, in contrast to MDN, DNp17 activity was not modulated during walking bouts, suggesting that DNp17 does not control walking parameters. In some cases, DNp17 appeared to be slightly inhibited during walking (Figure 3A), but overall, there was no correlation between DNp17 activity and walking parameters across walking bouts and flies (Figure 3B). This strongly suggests that DNp17 is not involved in the control of walking activity.

**Figure 3:**
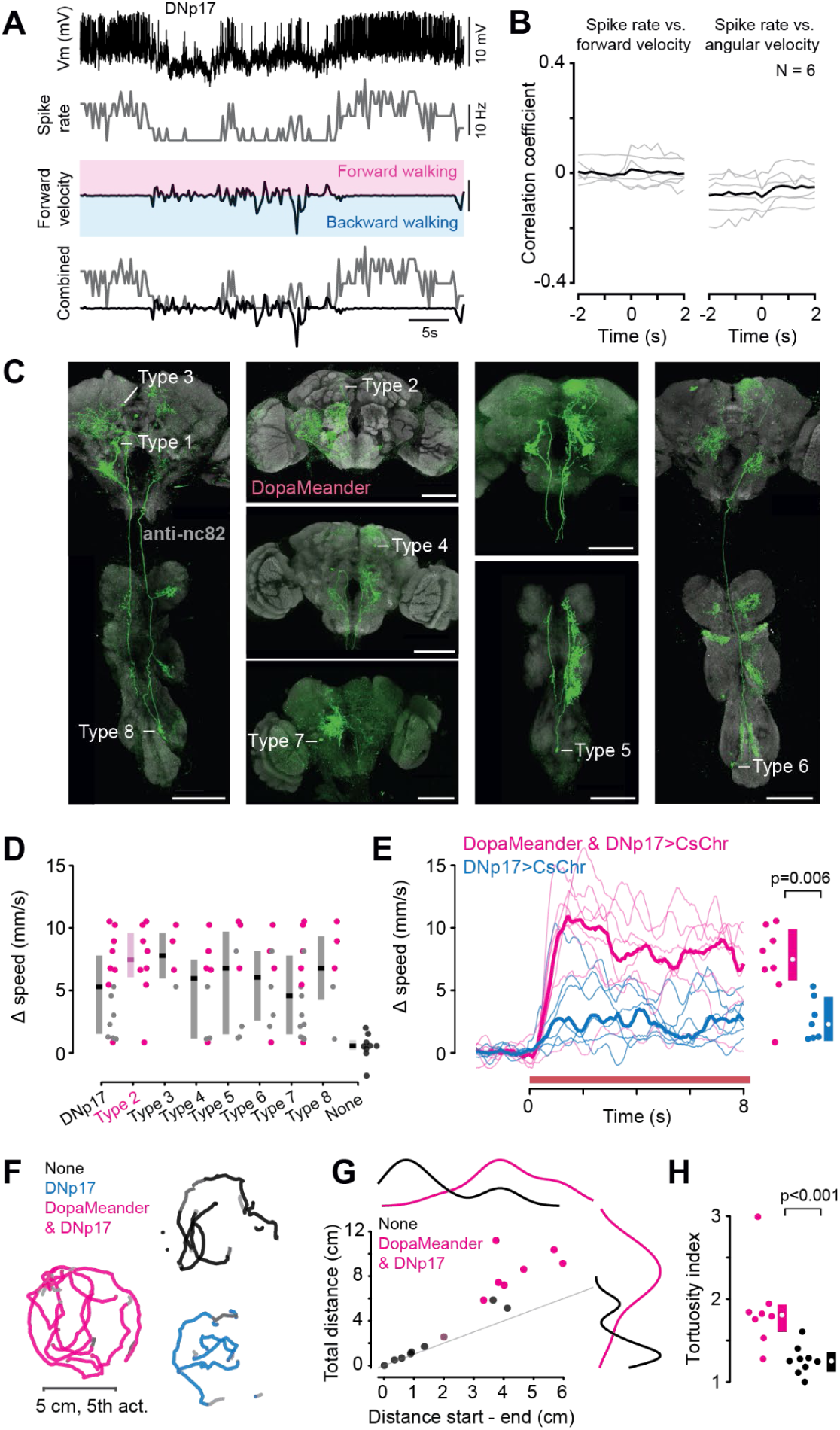
DNp17 activity is not correlated with walking parameters and the walking phenotype of the SS02553 line is best-explained by DopaMeander activation. A) Example recording of DNp17 membrane potential (Vm) during spontaneous walking, alongside the corresponding spike rate and forward velocity (both in 250 ms bins). B) Cross-correlation of DNp17 spike rate with forward and angular velocity reveals no correlation. C) Different neuron types labelled by the SS02553 line. CsChrimson::tdTomato was expressed stochastically in different neurons using SPARC2. In total, the SS02553 line labels 8 different neuron types. Neuron type 2 corresponds to DopaMeander (magenta). D) Average walking speed during optogenetic activation of individual flies in the UFO expressing SPARC2-CsChrimson in different subpopulations of SS02553. DopaMeander expression indicated in magenta. Flies expressing CsChrimson in other neuron types shown in gray, and flies without expression shown in black (‘none’). E) Increase in walking speed of flies expressing CsChrimson in DNp17 with type 2 (DopaMeander, magenta) was significantly faster than of flies without DopaMeander expression (blue). Red bar indicates optogenetic activation. F) Walking trajectories during optogenetic activation of flies expressing CsChrimson in no neurons (none), in DNp17 without DopaMeander, and in DopaMeander. Flies were tested individually, walking trajectories from different flies were overlaid for visualization. G) Walking activity of flies expressing CsChrimson in no neurons (black) and in DopaMeander (magenta). DopaMeander activation led to longer, more tortuous walking bouts. H) Tortuosity index for DopaMeander activation was significantly higher than for flies without expression in the CNS. Data for each fly was averaged across 9 trials at light intensity 4.1 mW/cm2 in all datasets. Boxplots show median and IQR in all panels. All p-values calculated via Wilcoxon rank sum test.

The DNp17 recordings strongly suggest that the forward walking and turning phenotype must be driven by other neurons labelled by the SS02553 line. To identify which of these neuron types were responsible for the walking phenotype, we used a stochastic labeling method (SPARC) to sparsely express CsChrimson:tdTomato in neuronal subpopulations targeted by the line^38^. This allowed us to optogenetically activate subsets of neurons in the SS02553 via CsChrimson and subsequently analyze which neurons were labeled via the tdTomato tag. To this end, individual flies were subjected to optogenetic stimulation in the UFO, after which their brain and VNC were dissected and imaged. We initially selected 18 flies which, based on our ad-hoc observation, displayed an increase in walking speed, and dissected them. This approach revealed that the SS02553 line targets seven different neuron types in addition to DNp17 (Figure 3C). This includes four local brain interneurons (type 2, 3, 4 and 7), two ascending neurons (type 5 and 8) and one local neuron in the VNC (type 6). These expression patterns were further confirmed via MultiColor FLPOut stainings^39^ (not shown) and comparisons to previously published images^10^.

This time-consuming method yielded a total of 26 flies that were used for behavioral analysis and subsequently stained and anatomically analyzed. Out of these, 17 flies expressed CsChrimson in several but not all of the neurons labelled by the SS02553 driver line. 9 flies showed no visible expression in any neuron. These flies exhibited very low average forward velocities, and hence, no walking phenotype (Figure 3D, ‘none’). Flies expressing CsChrimson in DNp17 displayed a broad range of forward velocities between 1 and 12 mm/s with a median of 5 mm/s. In contrast, all flies in which neuron type 2 was labeled exhibited a much higher forward velocity, independent of which other neurons were labelled alongside type 2. Three other neuron types (3, 5, and 8) drove increased forward velocities compared to flies without expression. However, this effect was driven by flies in which type 2 was also labeled. In contrast, flies in which type 2 was labelled without types 3, 5 or 8 still produced robust increases in forward velocity (Figure 3D). Hence, type 2 emerged as the strongest candidate for driving the forward walking phenotypes, as it consistently increased the walking velocity upon activation. Furthermore, flies with expression in type 2 and DNp17 walked significantly faster than flies in which DNp17 was labelled without type 2 (Figure 3E).

Since type 2 was responsible for the walking phenotype, and because it expresses dopamine (see below), we hereafter refer to neuron type 2 as ‘DopaMeander’. In addition to increasing forward walking, DopaMeander activation increased turning, as indicated by the raw trajectories, which were longer and more curved than those of DNp17-activated flies and flies without expression in the CNS (Figure 3F). We quantified this effect by measuring the distance and curvature of walking trajectories, as described previously. DopaMeander activated flies were consistently shifted towards the upper left corner of the behavioural space compared to flies without expression in the CNS (Figure 3G), resulting in significantly higher tortuosity indices for the population of DopaMeander activated flies (Figure 3H). The fine-grained behavioral and anatomical analysis based on the sparse SPARC-activation revealed that DopaMeander, and not DNp17, is responsible for the forward walking and turning phenotype driven by the SS02553 line. Based on these experiments, we hypothesised that DopaMeander plays a critical role in the neuronal control of forward walking and turning, and thus provides an interesting counterpoint to MDN, which controls backward walking.

### DopaMeander is a dopaminergic brain interneuron connected to locomotor circuits

Since DopaMeander was not previously described or characterized, we sought to shed light onto the neuron’s identity and connectivity. Using intracellular neurobiotin injections via patch-clamp recordings, we performed single-cell labeling of DopaMeander neurons to analyze their anatomy (Figure S3A). This enabled identification of a split-Gal4 driver line labeling DopaMeander in the FlyLight light microscopy dataset (Figure S3A) and matching DopaMeander to a unique cell type in the FlyWire electron microscopy (EM) volume^11,40^ (Figure 4A and S3A). Both matches indicate that DopaMeander is a dopaminergic neuron. The split-Gal4 line labeling DopaMeander is the dopamine-specific line SS45091^41^, which targets the TH and Ddc genes involved in dopamine synthesis. In the EM, DopaMeander is predicted to be dopaminergic based on neurotransmitter classification algorithms^42^.

**Figure 4:**
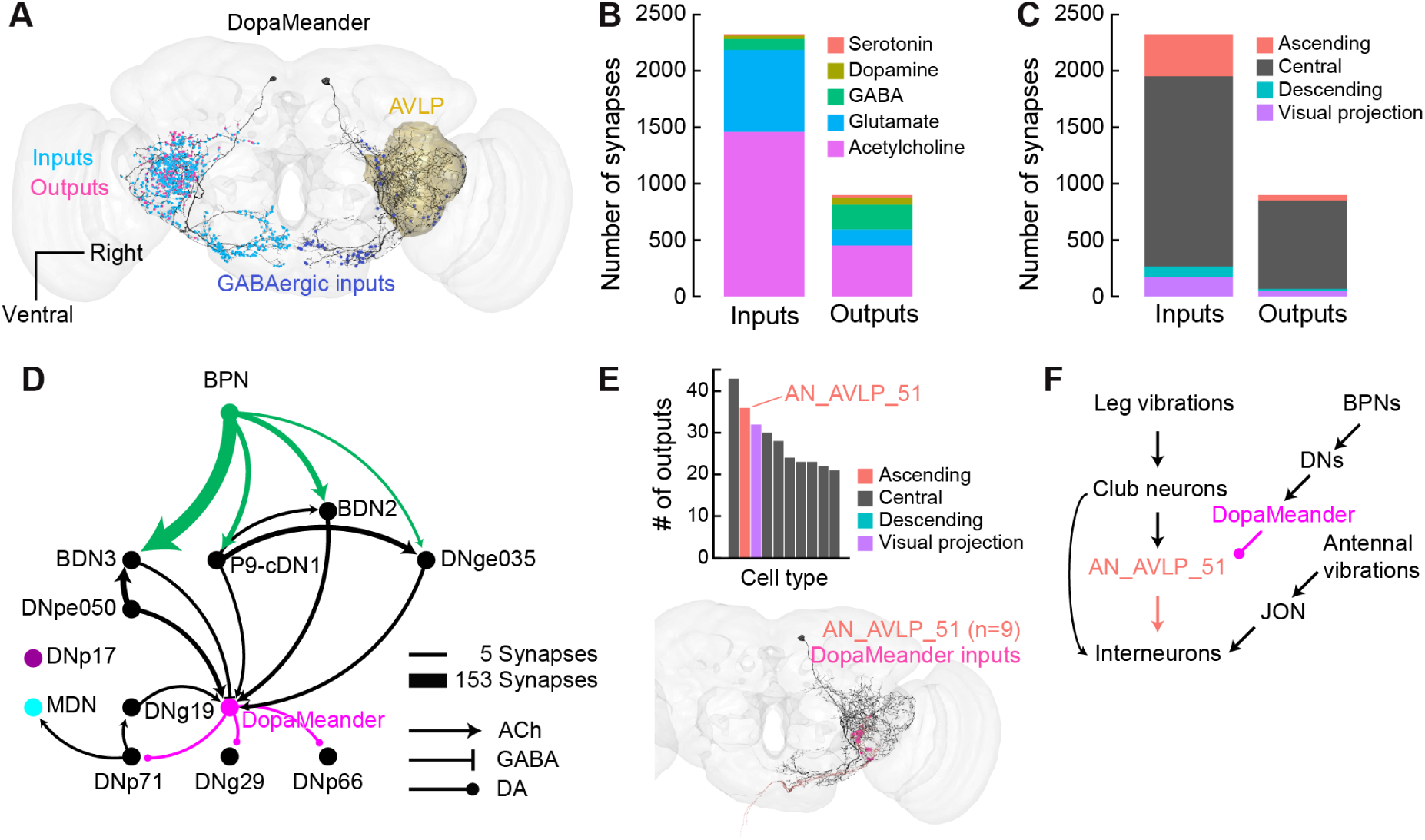
DopaMeander is connected to walking and leg mechanosensory circuits. A) EM reconstruction of DopaMeander, overlaid with presynaptic output sites (magenta), postsynaptic input sites (cyan), and GABAergic input sites (blue). The AVLP neuropil where the neuron has mixed polarity is highlighted in yellow. B) Number of total inputs and outputs of DopaMeander, colored by their predicted neurotransmitter^42^. C) Number of total inputs and outputs of DopaMeander, colored by neuron superclass^40^. D) Connectivity graph of DopaMeander with its descending partners (cyan superclass in (C)). DNp17, MDN and walking initiation BPN neurons^14^ are also included. Nodes represent cell types, edges depict synaptic connections. Edge color and thickness denote presynaptic neurons and synapse numbers. Edge shape depicts predicted primary neurotransmitter for each cell type. For clarity, edges with fewer than 5 synapses are not shown. E) Top, DopaMeander output distribution, grouped by cell type and colored by superclass. Only the ten strongest outputs are shown. The ascending population AN_AVLP_51 (salmon) receives strong axo-axonic input from DopaMeander. Bottom, EM reconstruction of AN_AVLP_51 neurons, overlaid with DopaMeander (black) and its synaptic inputs (magenta). F) Circuit diagram showing the mechanosensory circuit DopaMeander is embedded in. Leg vibration signals are transmitted to the brain via AN_AVLP_51 (identified as 8B/10B neurons in^44^). These signals are likely modulated by DopaMeander which relays walking-related signals from BPNs before they are integrated by downstream circuits that process antennal vibrations.

Using the match in the FlyWire connectome, we analyzed the anatomy and connectivity of DopaMeander in detail. DopaMeander neurons are a pair of local brain interneurons, projecting primarily to the anterior ventrolateral protocerebrum (AVLP) and to the gnathal ganglia (GNG) neuropils (Figure 4A). Both of these brain regions are implicated in the neuronal control of locomotion. The GNG serves as a central hub for the integration of descending and ascending neurons^40^, and is often compared to the vertebrate brainstem. The AVLP is particularly relevant for the integration of ascending feedback from locomotor networks^43^. DopaMeander input synapses stem primarily from cholinergic and glutamatergic neurons (Figure 4B), with a smaller number of GABAergic inputs concentrated in the GNG and near the primary neurite in the AVLP (Figure 4A).

Since DopaMeander activation induced forward walking and turning, we investigated if DopaMeander is connected to previously identified neurons involved in the control of walking. DopaMeander receives most of its inputs from central neurons innervating the AVLP, followed by several classes of ascending neurons from the VNC (Figure 4C, Figure S3C). While many of these inputs are from previously undescribed neurons, inputs from descending neurons originate from a subnetwork shared by Bolt Protocerebral Neurons (BPN), which are known to induce forward walking^14,23^ (Figure 4D). These DNs are downstream of BPNs and target DopaMeander but not DNp17, further suggesting that DopaMeander, and not DNp17, is embedded in a forward walking-related circuit. MDN, which promotes backward walking, is also not directly involved in this network (Figure 4D). Compared to its inputs, DopaMeander provides much fewer outputs, which generally consist of weak and broad synaptic connections to interneurons (Figure 4C, Figure S3D). This is a common motif among modulatory neurons, which presumably primarily act via volume transmission. Surprisingly, the second strongest output of DopaMeander are the axon terminals of AN_AVLP_51 (Figure 4E) – an ascending neuron type that receives about a third of its axo-axonic input from DopaMeander (Figure S3E). These ascending neurons belong to the 8B/10B hemilineage and are thought to relay leg vibration signals from the VNC to mechanosensory processing centers in the brain during walking^44^ (Figure 4F).

The meandering forward walking phenotype we observed could, in principle, arise from reciprocal inhibition between the left and right DopaMeander neurons. However, connections between the left and right DopaMeander neurons are polysynaptic and weak, and they are mediated largely via cholinergic and glutamatergic inputs, so that mutual inhibition is unlikely to be the source for the meandering phenotype (Figure S3B). The inhibitory GABAergic input both DopaMeander neurons receive in the GNG is therefore unlikely to balance left-right activity. Instead, it might contribute to bilateral gating of DopaMeander. In summary, our anatomical analysis and behavioral results strongly suggest that DopaMeander, and not DNp17, drives and encodes aspects of forward walking. DopaMeander is positioned within a walking-related circuit that processes ascending mechanosensory feedback from the legs, and descending mechanosensory signals from the antennae. Hence, it might play a role in the integration of internal states, via the modulatory component, and external sensory cues and ascending feedback, via the mechanosensory pathways.

### MDN and DopaMeander differentially encode walking parameters

Our behavioral results indicate that DopaMeander drives forward walking with a turning component, whereas the MDN population drives straight backward walking, in response to antennal contacts (see Figure 1) and visual stimulation^13^. If these two neuron types indeed control these opposite walking parameters in behaving animals, their activity patterns should differ significantly during spontaneous walking. To explore how DopaMeander and MDN contribute to the control of walking *in-vivo*, we recorded from both populations via patch-clamp in tethered flies during spontaneous walking over extended periods of time.

DopaMeander was spontaneously active and exhibited a spike rate between 0 and 40 Hz when flies were tethered and walking on the trackball (example in Figure 5A). DopaMeander was already active during non-walking periods, but strongly increased its spike rate during forward walking bouts (Figure 5A). MDN activity was also strongly modulated in tethered, spontaneously walking flies. In the example recording shown in Figure 5B, MDN spike frequencies ranged from 0 to 75 Hz. The strongest excitation of MDN occurred during bouts of spontaneous backward walking, during which MDN exhibited strong depolarizations and increases in spike rate. Conversely, MDN activity was attenuated during periods of resting, and was further reduced during forward walking bouts (Figure 5B). MDN and DopaMeander therefore contribute to centrally-generated, spontaneous forward and backward walking in the absence of deliberate sensory stimulation (see Figure 1).

**Figure 5:**
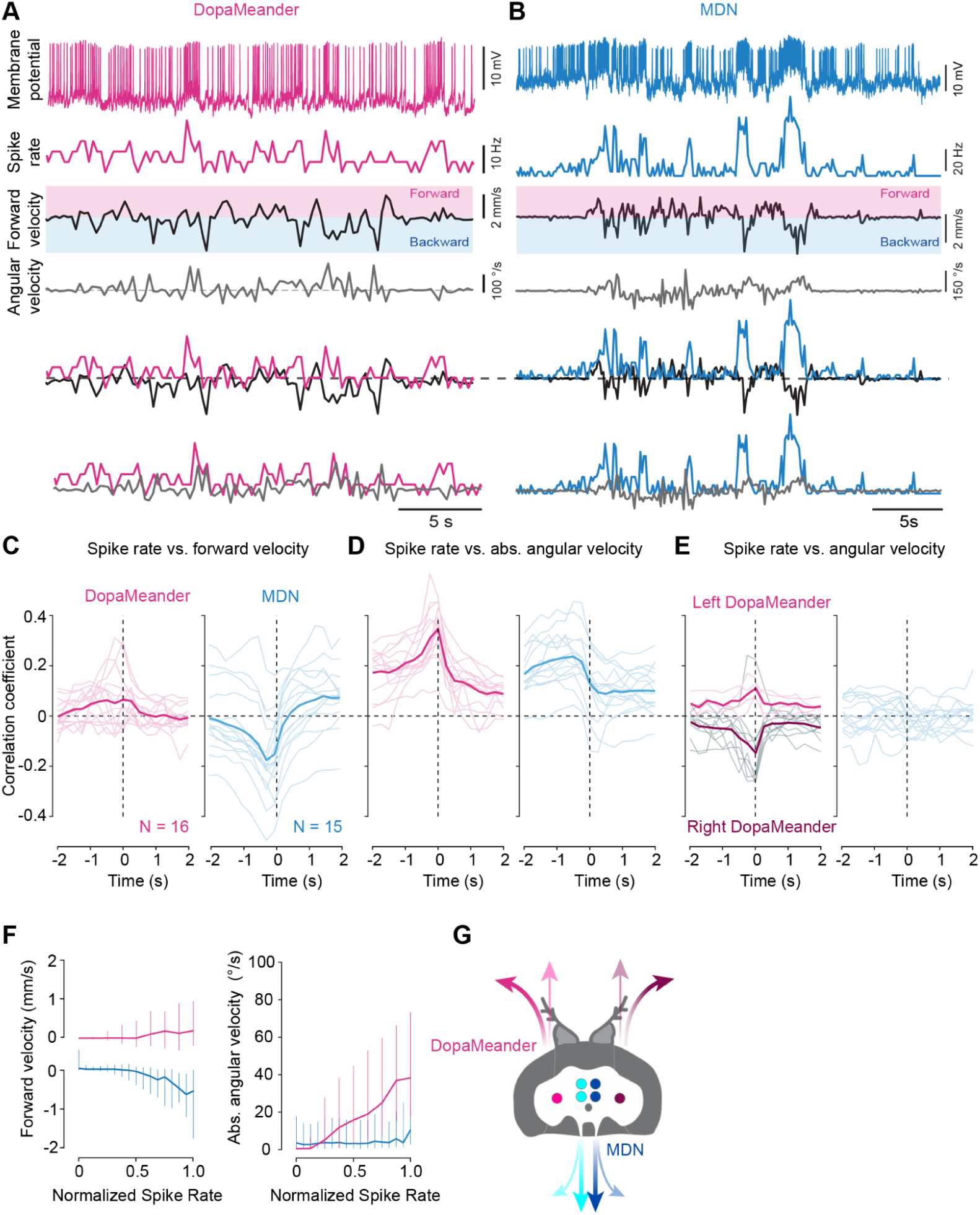
DopaMeander and MDN differentially contribute to backward walking, forward walking, and turning. A) From top to bottom: Example recording of DopaMeander membrane potential (Vm) during spontaneous walking, alongside the corresponding MDN spike rate (in 250 ms bins), and forward (magenta) and backward velocity (blue, both in 250 ms bins). Bottom traces, overlay of spike rate and forward/backward velocity and overlay of spike rate and angular velocity. B) Example recording of MDN during spontaneous walking. Plot details as in (A). C) Cross correlation analysis for DopaMeander (magenta, N=16) and MDN (blue, N=15) spike rates and backward velocity, both in 250 ms bins. All data sets were z-scored before correlation analysis. D) Same as (C) but for absolute angular velocity, i.e., turning. E) Same as (C) but for angular velocity, i.e., left or right turning. Left (light) and right DopaMeander (dark) populations could be distinguished during the recording. F) Normalized spike rate vs forward (left) and absolute angular velocity (right) for MDN (blue) and DopaMeander (magenta) during spontaneous walking. To account for the latency between neuronal activity and behavior, the binned spike rate and walking velocities were shifted to the peak of the cross-correlations in (C), i.e., by one 250 ms bin. G) DopaMeander activity is correlated with ipsiversive turning and to a lesser extent with forward walking. MDN activity is highly correlated with straight backward walking.

Next, we analyzed the contributions of DopaMeander and MDN to spontaneous walking in more detail. To this end, we cross-correlated the spike rate and membrane potential of DopaMeander and MDN with the translational (forward vs. backward) and rotational (turning) components of the walking trajectories (Figure 5C-E, see Figure S4 for membrane potential correlations). Increases in both DopaMeander and MDN activity typically preceded walking, as is evident in the shift of the cross-correlation peaks to the left (Figure 5C). More specifically, DopaMeander was weakly but positively correlated with forward walking, while MDN was negatively correlated with forward walking. Hence, DopaMeander activity increases slightly during forward walking, whereas MDN activity increases strongly during backward walking – in line with the activation phenotypes of the two neurons. In contrast to the weak correlation with forward walking, DopaMeander activity was strongly correlated with the absolute angular velocity, and thus turning (Figure 5D). MDN, on the other hand, reduced its activity during turning, in line with a primary role in straight backward walking.

In the case of DopaMeander, we could easily determine whether we recorded from the left or the right neuron based on its soma location, which was not possible for MDN since all four somata of the MDN population lie along the midline. When splitting the DopaMeander population into left and right neurons, we found a lateralized correlation between spike rate and angular velocity. Higher spike rates in the right DopaMeander were linked to increased angular velocity, and hence turning towards the right, and vice versa for the left DopaMeander (Figure 5E). This suggests that DopaMeander activity is lateralized, and that each neuron facilitates turning towards the ipsilateral side. This activity pattern makes sense in light of the activation phenotype, in which activation of DopaMeander drove walking with increased turning.

Both populations, MDN and DopaMeander, maintained a consistent tonic spike rate during rest phases, where no leg movements occurred. This constant activity provides a means to control walking direction by either elevating or suppressing neuronal activity, offering a higher dynamic range for controlling walking compared to a simplistic on/off switch. Supporting this idea, both populations exhibited an activity tuning curve, with higher neuronal activity occurring during faster backward walking and turning, respectively (Figure 5F). In summary, MDN activity was strongly correlated with backward walking but not turning, whereas DopaMeander activity was weakly correlated with forward walking and strongly correlated with turning, the latter with an ipsiversive bias. These correlations are in line with their respective activation and silencing phenotypes (Figure 5G).

### MDN and DopaMeander control opposite features of spontaneous walking

Our correlation analysis revealed that MDN and DopaMeander activity is linked to straight backward walking and forward turning, respectively. However, a cross-correlation does not quantitatively describe how neuronal activity is dynamically translated into walking behavior. To address this, we used linear-nonlinear (LN) models to predict behavioral components from MDN and DopaMeander activity during spontaneous walking (Figure 6A, B). The LN model consists of a linear temporal filter of the neuronal activity and a nonlinearity that maps this activity to walking behavior, and thus motor output. This approach provides an interpretable framework for understanding how MDN and DopaMeander control walking by capturing both the temporal structure of neuronal activity and the nonlinear transformation to behavior. We decomposed different walking parameters from the treadmill rotation, and evaluated how much of the variance in unprocessed (raw), positive, negative, and absolute components of translational (forward/backward) and angular velocities could be explained by each population’s activity (Table S2), and the estimated linear filter of this computation.

**Figure 6.**
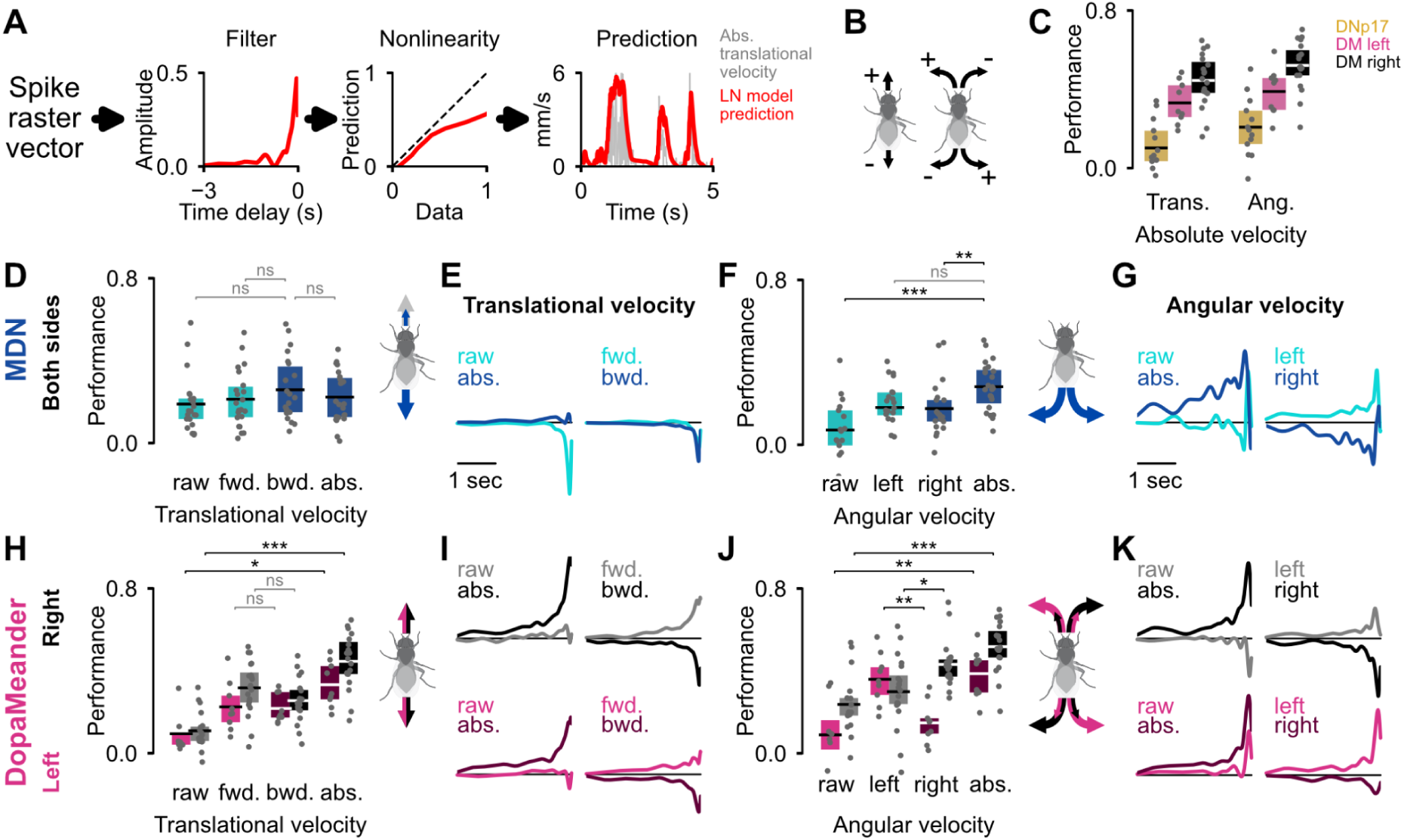
MDN and DopaMeander activity predict distinct components of locomotion during walking. A) Schematic of the linear-nonlinear (LN) model. Left to right: example filter, nonlinearity, and prediction for a 5-second segment of absolute translational velocity from a model trained with DopaMeander activity. B) Schematic of velocity components and sign conventions. C) Comparison of performance on the held-out data of models fit to translational and angular absolute velocity, for DNp17 (gold) and DopaMeander (left: magenta, right: black) recordings. D) Performance of MDN translational velocity models. E) Average filters from the MDN models fit to components of the translational velocity. Left: raw and absolute values; right: forward (fwd.) and backward (bwd.) components. F) Performance of MDN angular velocity models. G) Average filters from MDN angular velocity models. H) Performance of DopaMeander translational velocity models. I) Average filters from DopaMeander translational velocity models. Top: right side neurons in black; bottom: left side neurons in magenta. J) Performance of DopaMeander angular velocity models. K) Average filters from DopaMeander angular velocity models. Individual points in (C), (D), (F), (H) and (J) represent individual recordings of the corresponding neuron activity, and the boxplots show mean and IQR. Lighter shades are used for raw and positive (forward or left) components to differentiate them from absolute and negative (backward or right) components in darker shade. Illustrations in (D), (F), (H) and (J) summarize the results. For all pairwise comparisons (in (D), (F), (H), (J)), we used Shapiro-Wilk to assess normality, applied t-tests or Mann-Whitney U tests accordingly (see Methods), and used Bonferroni-corrected p-values. Significance levels are as follows: *:<0.05, **:<0.01, and ***:<0.001. See Table S3 for statistical tests and explicit p-values per comparison.

First, we wanted to quantify any potential effects of DNp17 on walking, which we might have missed in our classical analysis approach (Figure 3A, B). As expected, models trained with DNp17 activity showed poor performance across all walking parameters (Figure 6C, Figure S5A), confirming that DopaMeander, and not DNp17, is the behaviorally relevant population targeted by the SS02553 driver line (Figure 3).

LN models trained on MDN activity predicted all translational locomotion components similarly well, without significantly better performance for backward vs. forward walking (Figure 6D, S5C). However, the filters estimated from forward, backward, and absolute component models are all negative (Figure 6E), suggesting that MDN activity explains increased backward walking and decreased forward walking. This implies that MDNs do not simply induce backward walking, but more generally contribute a backward-directed component to the translational velocity. Therefore, MDN activity can slow down forward walking up to the point that it reverses walking direction.

MDN activity best predicts absolute angular velocity, and thus the amount but not the direction of turning (Figure 6F). The estimated filters were broader for rotational velocities (Figure 6G), implying that MDN spike rate affected turning over longer time windows. The filters for left and right components were positive and negative, respectively, indicating that MDN activity predicted increases in angular speed in either direction (Figure 6G, S5D). Importantly, MDN activity predicts turning speed during backward walking, but not during forward walking (Figure S5E).

In contrast, DopaMeander activity predicts overall locomotor drive (Figure 6H). The filters for absolute translational velocity were positive, while the filters for forward and backward components were correspondingly positive and negative, indicating that DopaMeander activity increases walking speed in both forward and backward directions (Figure 6I, S5F).

Consistent with this, DopaMeander activity also best explains the speed and direction of turning, with a strong ipsiversive bias (Figure 6J-K), suggesting that the left DopaMeander drives left turns (positive) and the right DopaMeander drives right turns (negative). Interestingly, both hemispheres contributed to the prediction of contraversive turns, albeit with lower performance, indicating correlated bilateral activity during turning (Figure 6J, S5G). This is consistent with neural activity correlations, where DopaMeander spike rate is strongly correlated with overall turning (Figure 5D) while maintaining an ipsiversive bias (Figure 5E). Moreover, unlike MDN, DopaMeander predicts turning speed during both forward and backward walking (Figure S5H), emphasizing its role in modulating overall turning strength, not just during forward walking.

In summary, our modeling results show that the activity of MDN and DopaMeander predicts distinct locomotor features: MDNs are not only tuned to backward velocity and associated turning, but also to negative changes in translational velocity. DopaMeander neurons encode the overall walking gain and ipsiversive turns.

### Neurons controlling walking are gated out during flight

Brain dynamics are heavily affected by internal state changes^45,46^. In insects and vertebrates, behavioral state changes have particularly strong effects on sensorimotor pathways. In the cases we are aware of, the activity of visual inter- and descending neurons in *Drosophila* and other insects is boosted during flight^47–53^. These changes are thought to support the particular sensorimotor requirements of flight, for example faster visual processing, steering, and landing. MDN is firmly established as a walking-related DN, and its spike rate is directly correlated with walking. DopaMeander is a dopaminergic modulatory neuron, whose activity suggests that it is involved in the control of forward walking and turning. Hence, these two neuron types are components of the neuronal networks controlling walking, but operate on different levels: MDN connects visual interneurons in the brain’s periphery with motor networks in the VNC, while DopaMeander is positioned to modulate walking-related networks in the brain. Using these neurons, we can therefore ask how distributed networks supporting walking are affected by behavioral state changes into and out of flight. To test this, we employed an experimental setup allowing for patch-clamp recordings while flies transition between walking and flight, achieved by lowering the walking treadmill and thus removing ground contact^54^.

When recording from MDN, we noticed a steep hyperpolarization of the membrane potential upon removal of the treadmill, when the fly lost ground contact, followed by a further hyperpolarization step as flies initiated flight (Figure 7A). The spike rate also dropped markedly after flight onset and was reduced from 13.8 Hz before flight to 0.3 Hz during flight (Figure 7B). Throughout the flight bout, MDN maintained strongly hyperpolarized (−8.4 mV vs. pre-flight potential), with a slight repolarization before cessation of flight (−3.8 mV, Figure 7B). The spike rate was very low throughout flight, under 0.7 Hz on average, and gradually returned to baseline levels post-flight (Figure 7A, B). Hence, MDN was basically quiescent during flight periods, and its activity was much lower during flight than during non-walking periods on the treadmill.

**Figure 7:**
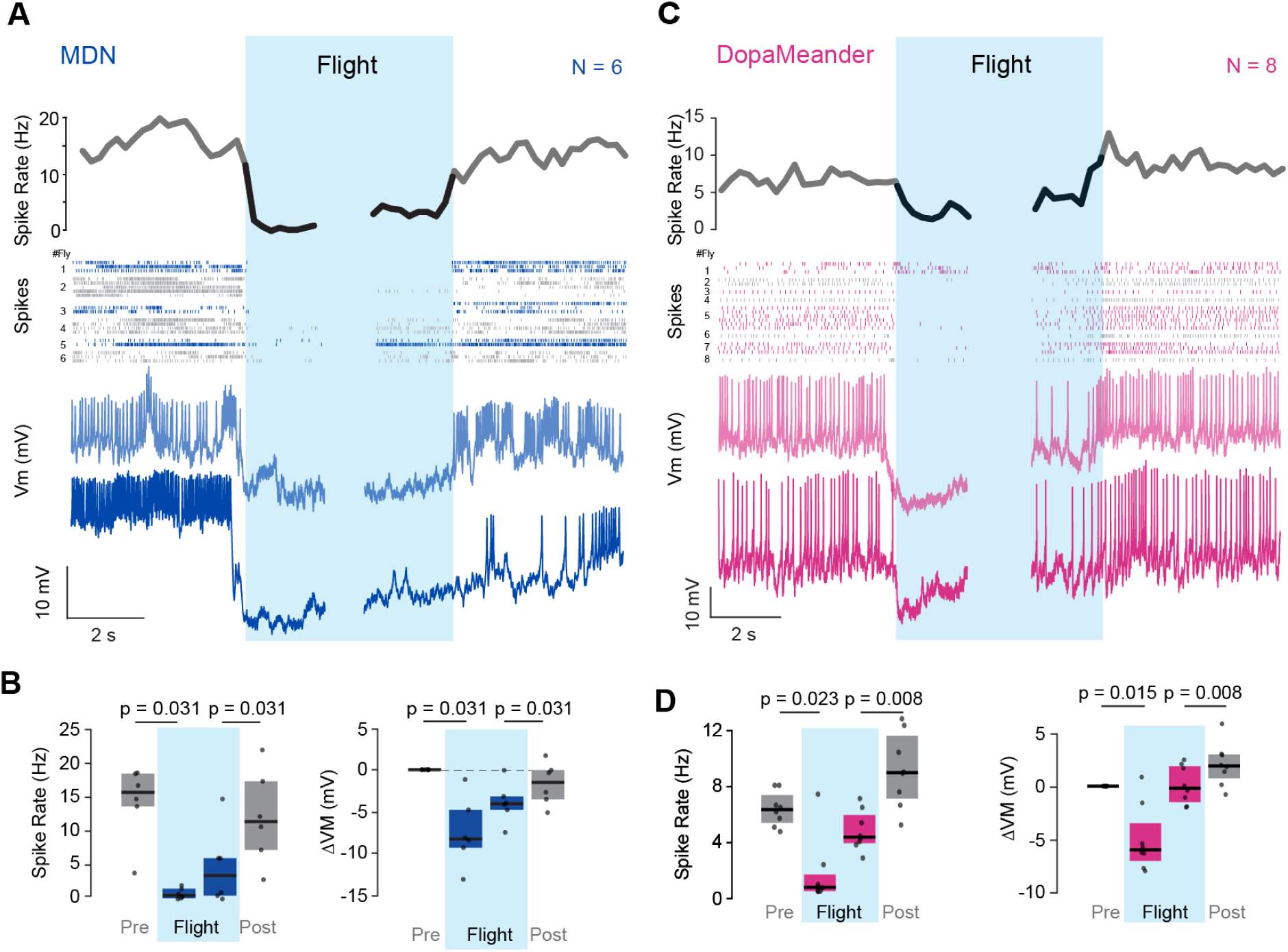
MDN and DopaMeander are gated out during flight. A) MDN activity before, during (blue shading) and after flight. Top, mean MDN spike rate (in 250 ms bins) across all trials and flies. Middle, spike events from all trials (colors alternating between flies). Bottom, membrane potential of two example recordings. B) Quantitative analysis of spike rates (left) and membrane potential (right) of MDN before (Pre), during, and after (Post) flight. Spike rates were analyzed in four 2 s long analysis windows: Pre-onset of flight, post-onset of flight, pre-cessation of flight, and post-cessation of flight. The membrane potential was analyzed in four analogous 1 s windows. The pre-flight potential was subtracted from subsequent windows. Data is shown as median ± IQR with individual data points representing different neurons recording in different flies. P-values were calculated using the Wilcoxon signed-rank test. C & D) Same analysis and plot details as in (A & B), but for DopaMeander activity.

Next, we performed the same experiments with DopaMeander. Similar to MDN, DopaMeander was immediately and strongly hyperpolarized at flight onset. The membrane potential was hyperpolarized by an average of −5.7 mV in the first two seconds of flight and maintained hyperpolarized throughout the flight duration. The membrane potential, however, was significantly repolarized during the final seconds of flight compared to the first seconds (−5.5 mV vs −0.6 mV; Figure 7D, right). The spike rate basically followed the membrane potential, and dropped to about 0 Hz in the first two seconds of flight, but then gradually increased until it reached 3.6 Hz before flight cessation (which occurred spontaneously). After cessation of flight, the spike rate of DopaMeander was slightly but significantly higher than before flight (9.7 Hz vs. 7.8 Hz; p = 0.004; Figure 7D, left).

These observations demonstrate that both MDN and DopaMeander undergo a strong hyperpolarization and exhibit reduced activity during flight, a behavioral state in which their contribution to motor control is presumably detrimental. Despite both populations being similarly affected by flight, they exhibited differences in the dynamics. DopaMeander was less hyperpolarized and more active towards the end of flight, whereas MDN stayed hyperpolarized throughout the whole flight duration. Hence, the effect of behavioral state was stronger and longer lasting for MDN. This might indicate that the two populations are gated by different mechanisms. Nonetheless, the overarching finding is clear: both walking-associated populations are essentially switched-off during flight, highlighting that state-dependent gating occurs across motor control hierarchies. Moreover, the results show that the gating is bidirectional, in that walking-related populations are gated out during flight, whereas flight-related populations are gated out, or less active, during non-flight periods^47–53,55^.

## Discussion

### MDN is a command-like neuron that integrates antennal mechanosensory and visual stimuli to drive backward walking

In this paper, we investigated two neuronal populations which modulate walking direction in *Drosophila*. The first, MDN, reverses walking direction from forward to backward. It was previously shown that MDN receives visual, mechanosensory, and olfactory input to drive backward walking^13,30,56^. Here, we present the first *in-vivo* recordings from MDN in spontaneously walking flies. Our recordings revealed that MDN is active during spontaneous backward walking in the absence of specific sensory stimulation. Hence, the MDN population fulfills all criteria for being command-like neurons for backward walking.

Moreover, we show that MDN integrates inputs from the ipsi- and contralateral antenna to initiate backward walking. Notably, MDN responses were phasic during antennal stimulation, which matched the brief nature of backward walking bouts, and MDN only responded to inward deflection of the antennae – there was no response during the return movement upon antennal release. Hence, MDN is tuned to the antennal movements which readily occur when the fly walks into an obstacle, a conspecific, or a predator. Interestingly, MDN was previously shown to contribute to backward curve walking during activation of upstream visual projection neurons, and individual MDNs can drive turning upon unilateral optogenetic activation^27,29,30^. Our results indicate that the MDN population drives straight backward walking in response to unilateral antennal stimulation. These results suggest that the MDN population can either be activated bilaterally, by antennal inputs, to drive straight walking, or unilaterally, by specific visual inputs, in which case they contribute to backward turning^13^. Consistent with this, our array of bilateral wide-field visual motion stimuli did not induce direction-selective responses, suggesting unilateral MDN activation might be primarily driven by unilateral small-field stimuli, like an ipsilateral visual loom, which excites upstream LC16 neurons. Interestingly, MDN silencing during optogenetic activation of LC16 visual inputs also had much stronger effects on the translational than the rotational component. I.e., flies with silenced MDNs walked significantly less backwards, but still turned during LC16 activation, underlining the tendency for the MDN population to drive straight backward walking^13^.

Our silencing experiments also revealed that flies without active MDNs still walked backwards in response to antennal touch, albeit at lower speeds and for shorter durations. Moreover, silencing affected backward walking more than backward turning. This suggests some degree of redundancy, and the existence of parallel descending pathways involved in the neuronal control of backward walking and turning. Previously, DNs which integrate antennal touch or movement and project to VNC motor networks were identified in cockroaches, crickets, stick insects, and other insect species^17,19,57–61^. Hence, descending antennal mechanosensory pathways for backward walking might be evolutionarily conserved within insects, and serve the general purpose of reversing direction in response to antennal touch.

One surprising finding was that MDN was still active during slow forward walking and during periods of rest, which was particularly clear from the LN model predictions. Essentially, MDN exhibited an increasing firing rate from slow forward walking to stopping to backward walking, during which it was predictive of backward walking speed. This suggests that, rather than being exclusively recruited during backward walking, the MDN population could also serve to slow down forward walking and facilitate the transition to stopping and, finally, a reverse in walking direction.

Currently, several mechanisms to slow down walking are proposed, e.g. the inhibition of DNs that drive forward walking within the brain or the activation of neurons in the VNC that lead to an active arrest of stepping movements^23^. MDN connectivity in the larva, in which MDNs have the same function as in the adult and drive backward crawling, revealed multi-hop inhibitory connections from MDN to neurons driving forward walking^30^. Hence, MDN connectivity in the larva suggests they could contribute to the termination or reduction of forward walking.

Populations of neurons that stop or slow down walking were also identified in the vertebrate CNS, suggesting that the organization of the neuronal control of locomotion with specific populations for different parameters, i.e., starting, maintenance, stopping, and turning, is a shared principle of motor control^62–66^. Our results support the idea that walking is controlled by different populations contributing competing drives, such that, in case of backward walking, flies eventually switch walking direction if MDN activity supersedes the forward walking drive provided by other populations, including DopaMeander. Consistent with this, flies occasionally walk forward during long-term activation of MDN, suggesting forward walking is not completely suppressed by MDN^23^. The fact that MDN was not responsible for backward turning in antennal stimulation experiments suggests that MDN is not the only DN contributing to the backward walking drive. This is also supported by the observation that olfactory-driven backward walking is partially MDN independent^56^.

Additionally, components of the stepping generating networks in the VNC, such as levator or depressor muscles of the legs, serve the same function during locomotion independent of the walking direction, as demonstrated in cats^67^. MDN could contribute to the drive these muscles receive during both forward and backward walking.

### DNp17 is unlikely to drive fast forward walking

We confirmed previous data showing that the SS02553 split-Gal4 driver targeting DNp17 increased walking speed^22^. While many of the DN lines generated by Namiki et al. are extremely sparse and specific for the DN targeted, the SS02553 line labels several additional populations of neurons in the brain and VNC^10,68^. We confirmed that DNp17 is unlikely to be responsible for the forward walking activation phenotype in two ways: First, we showed that DNp17 activity is not correlated with forward walking via patch-clamp recordings in behaving flies. Second, we used a stochastic labeling approach to show that DNp17 increases walking speed only slightly, if at all, but that DopaMeander labeling is much more predictive of forward walking. This is also in line with DNp17 does not project to leg-related neuropils in the VNC on light or EM levels. In fact, DNp17 mainly targets flight related neuromeres such as the lower tectulum, haltere-related neuropils in the tectulum, and the neuropils controlling neck movement in the tectulum, with very few outputs in the leg neuropils^10^. Moreover, in contrast to many other DNs, DNp17 does not provide extensive outputs to neuropils in the gnathal ganglion, which serves as an integration center for walking circuits^10,69,70^. Based on its distribution of output synapses, the lack of a clear activation phenotype during walking, and the fact that DNp17 is not correlated with walking activity, we strongly suspect that DNp17 is more relevant for flight control than for walking.

### DopaMeander is a central dopaminergic neuron that contributes to forward walking and turning

Three lines of evidence suggest that DopaMeander contributes to the control of forward walking and turning. First, DopaMeander labeling was the best predictor of fast forward walking in stochastic activation experiments (Figure 3D, E). Our initial data set focused on flies which seemed to increase their walking speed during activation, which were subsequently dissected. In 8 out of 26 flies tested, DopaMeander was labeled. The presence of additional neurons labeled in individual flies did not make a phenotype more or less likely, strongly suggesting that DopaMeander drives the fast walking phenotype. Second, DopaMeander activity correlates with forward walking, but even more strongly with turning, with an ipsiversive bias. This matches the phenotype of activating the driver line. Third and finally, connectome analysis revealed that DopaMeander connects to neurons involved in walking control and leg mechanosensation. Care needs to be taken in interpreting this connectivity. DopaMeander is a dopaminergic modulatory neuron, which is also evident in its basic connectivity pattern because it receives many more input synapses than it provides outputs. Hence, DopaMeander likely exerts a large proportion of its effects via volume transmission, which would not be visible in the connectome. This is common for dopaminergic neurons, as evident for instance by the fact that dopaminergic neurons in the mushroom body (DANs) make very few synapses in direct vicinity to Kenyon-Cell – mushroom-body-output-neuron (MBON) synapses^71^, but are still capable of modulating the synaptic strength of this connection significantly during learning^72,73^.

However, the sparse output synapses DopaMeander does provide to downstream neurons are in line with a role in locomotion. Moreover, DopaMeander receives and delivers a large proportion of its synaptic input and output in the anterior ventrolateral protocerebrum (AVLP) – a brain region closely associated with integrating sensory input with ascending feedback from VNC leg motor centers, which receives strong ascending input correlated with walking^43^. It is possible that each DopaMeander, when active, releases dopamine into the ipsilateral AVLP and thus boosts activity unilaterally to facilitate walking and turning, in particular in the ipsiversive direction. While it is tempting to speculate on the conserved role of dopamine in motor control and arousal across insects and vertebrates^74,75^, we did not investigate the role of dopamine itself for the function of DopaMeander, or the dopamine receptor distribution in the AVLP, so that we cannot make strong statements in this direction.

Our DopaMeander and MDN recordings also provided interesting insights into asymmetries in the control of walking, and hence turning. MDN is clearly capable of driving turning due to asymmetric outputs in the VNC by each MDN^13,28^. However, our results reveal that this asymmetry is compensated for by each MDN receiving bilateral input from the antenna, leading to straight backward walking in this context. The same seems to apply to visual inputs, since we found no directional preferences for each MDN in the population. Hence, MDNs seem to respond to stimuli from both directions equally well, leading to symmetric drive to the VNC. This is, for example, also the case in the giant fiber escape system, which receives strongly lateralized visual input on the connectome level^76^, but displays bilateral visual responses^77^.

DopaMeander, on the other hand, drove increased turning even during bilateral activation. Since we did not find any evidence for direct, inhibitory connections between the left and right DopaMeander, which might have been responsible for the symmetry break, the pathways downstream of DopaMeander must drive asymmetries in activity.

### Neurons controlling walking across hierarchical levels are gated out during flight

By recording from MDN and DopaMeander during behavioral state transitions, we were able to show that both neuron types are hyperpolarized and reduce their spike rate strongly and significantly during flight. Hence, they are gated out. In the motor control hierarchy, MDN and DopaMeander take opposite spots in the brain circuitry. DopaMeander is a modulatory neuron that likely exerts its effects through the modulation of a broad population of neurons in a brain region that integrates ascending feedback with central drive to control walking. MDN, on the other hand, leaves the brain and directly innervates motor circuits in the VNC. The fact that both neurons are gated out during flight suggests that walking-related neurons are inhibited during flight across layers of the motor control hierarchy. This in turn suggests that there are dedicated sets of neurons for the control of walking and flight across hierarchical levels. Earlier studies have shown that octopaminergic neurons exhibit the opposite state-dependence and increase their activity during flight^78^. Insulin-Producing Cells, on the other hand, are strongly inhibited during flight periods^54^. Together, these results highlight that different modulatory systems differentially shape sensorimotor circuit activity in an orchestrated and state-dependent way. Presumably, this also applies to space – modulatory neurons innervating different brain regions and compartments are likely modulated independently. For instance, different types of dopaminergic neurons in the mushroom body of the fly are differentially modulated during locomotion in the context of odour navigation^79^. Similarly, on the DN layer, neurons controlling the leg movements that occur during landing responses in flying flies are only active during flight bouts and unresponsive to visual stimuli during non-flight periods^50^. The same applies to other populations of flight-related DNs^52,53,55^. Our MDN gating result expands the concept of DN gating. So far, most studies were consistent with the idea that DN population activity, like neurons in the visual system, are generally boosted during flight periods. The fact that MDN, as a walking-specific DN, exhibits the opposite state-dependence strongly suggests that DN ensembles are gated in and out according to the behavioral requirements of the animal in different states, such that the flight control systems are only accessible to the brain during flight bouts, whereas walking-related motor systems are only accessible while the legs have ground contact. This likely serves as a mechanism to avoid maladaptive behavioral crosstalk, such as generating walking-like movements during flight in response to headwind.

How this gating is achieved mechanistically is still unclear. The fact that DopaMeander was consistently less hyperpolarized during later stages of flight whereas MDN stayed strongly hyperpolarized throughout flight suggests that two independent mechanisms could be implemented. Even neurons controlling the same motor behavior on the same level – for instance, two DN types controlling landing – can use separate mechanisms for state-dependent gating^50^. Our results suggest multiple parallel mechanisms are also used to modulate neurons across hierarchical levels.

## Key Resources Table

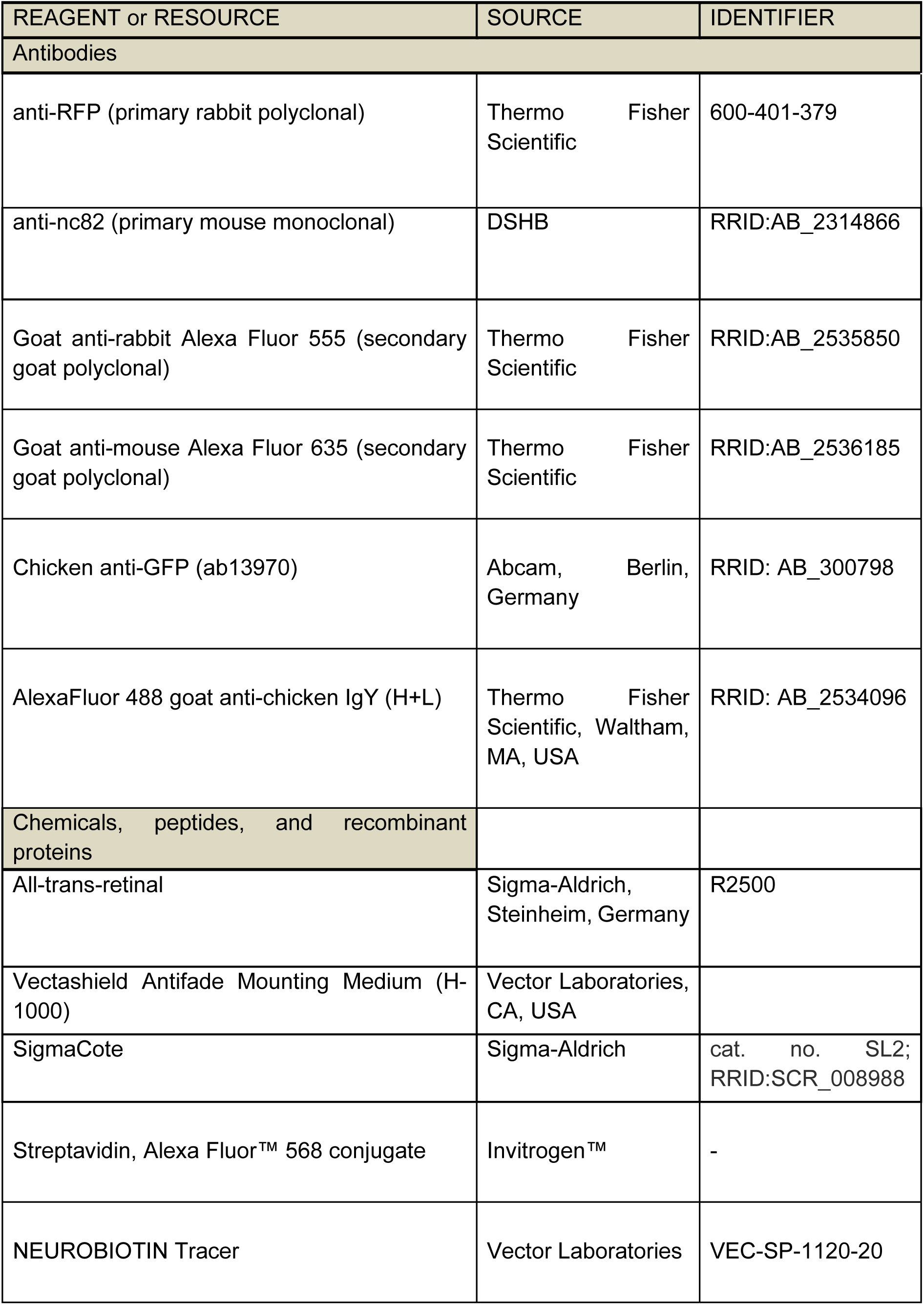
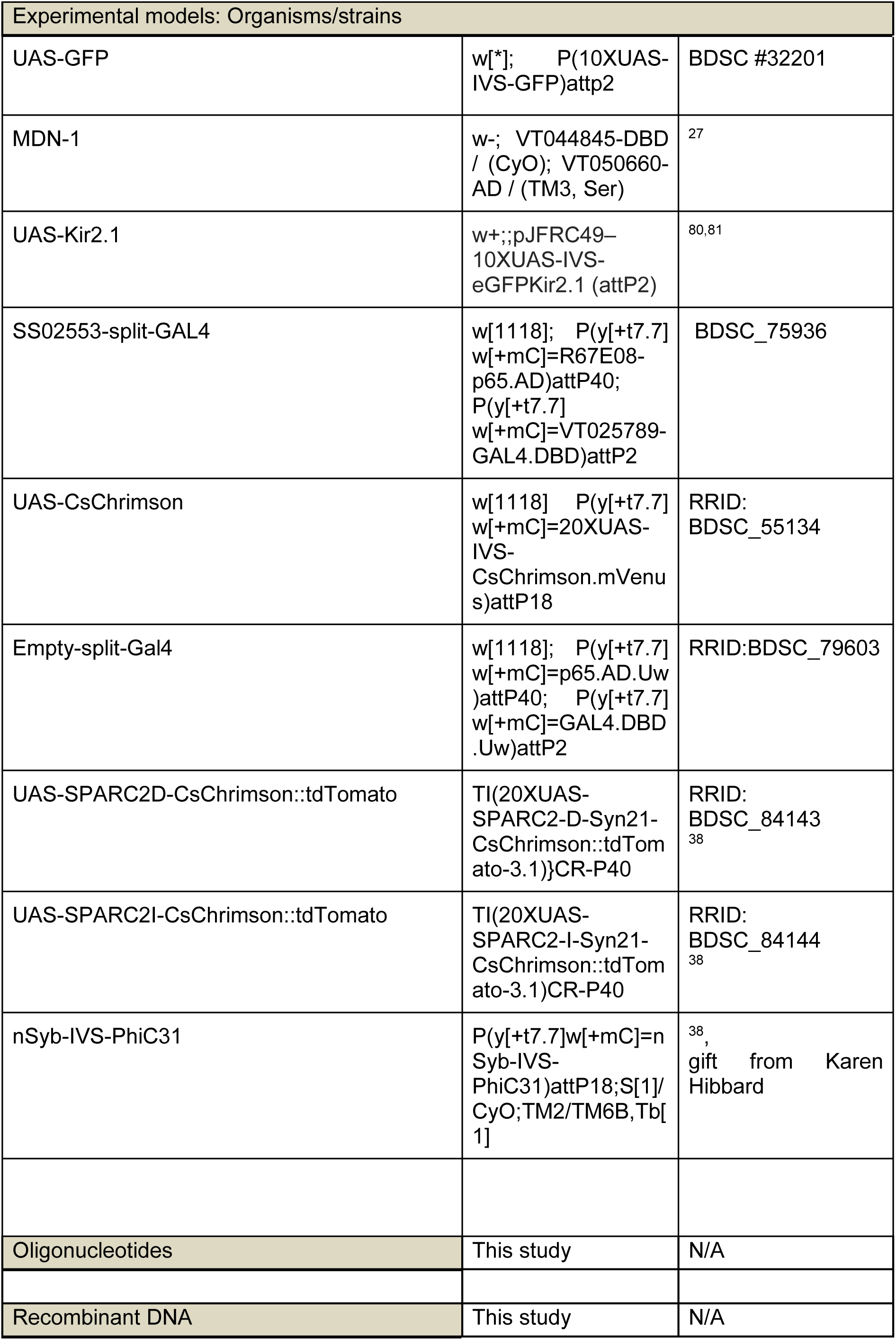
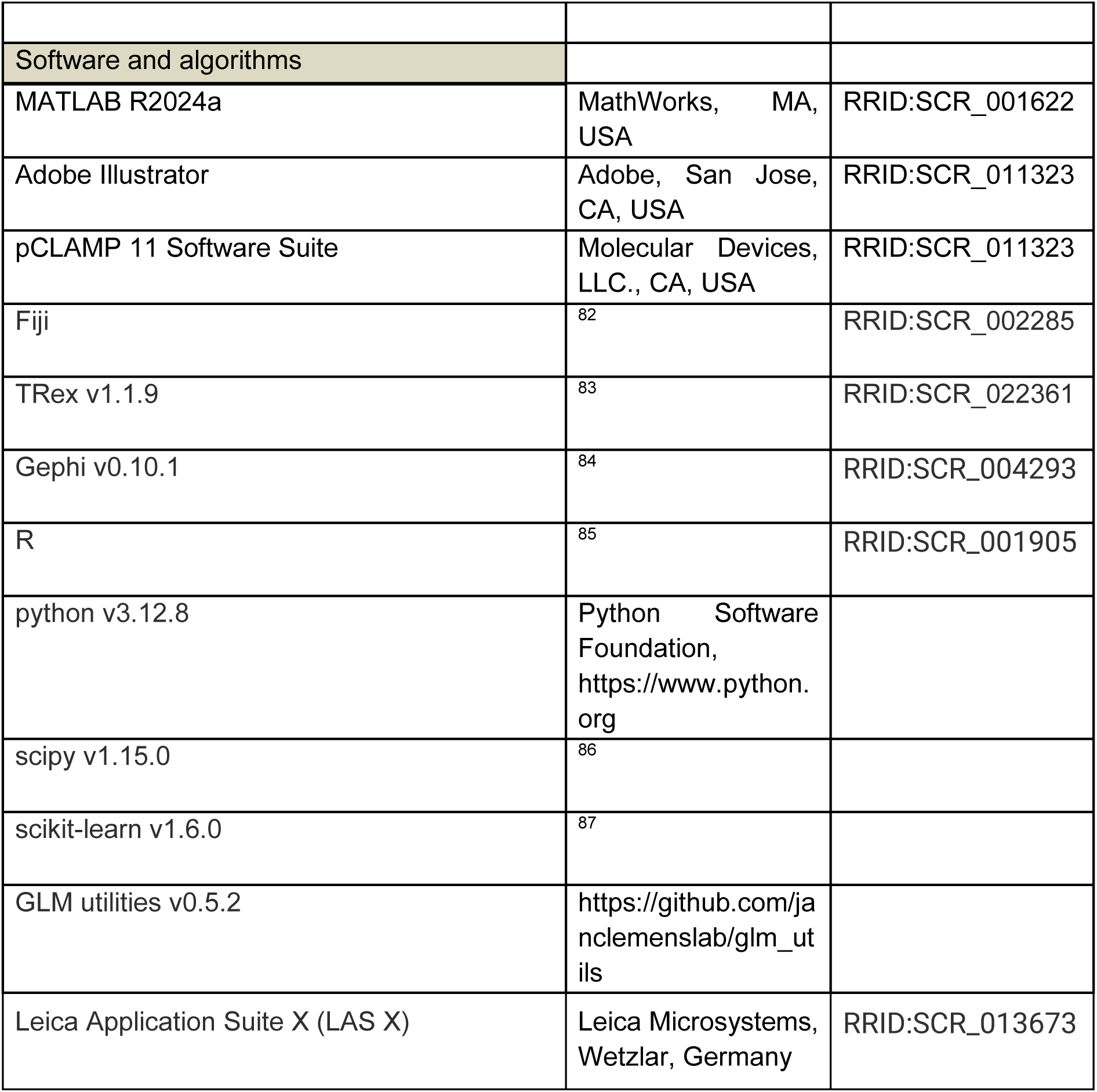

## Methods

### Resource availability

#### Lead Contact

Further information and requests for resources and reagents should be directed to and will be fulfilled by the lead contact, Jan M. Ache.

### Experimental model and subject details

#### Fly husbandry

All *Drosophila melanogaster* stocks were reared on standard fly food (cornmeal-agar-molasse medium) and maintained at 25°C and 60% relative humidity under a 12h/12h light/dark cycle. All electrophysiological experiments were performed on mated female flies 3-6 days after eclosion. For optogenetic activation experiments, post eclosion equal number of male and female flies were reared on standard fly food coated with 300 µM all-trans-retinal (R2500, Sigma-Aldrich, Steinheim, Germany) for a period of 3-6 days prior to the experiment. The flies were reared and maintained in the dark until used for the experiment.

#### Fly stocks and genotypes

The genotypes used for each figure and dataset are described in Table S1 and are listed in the key resources table.

## Method Details

### Electrophysiology

For *in-vivo* patch-clamp experiments, flies were handled and prepared as described before^54^. For *in-vivo* whole cell patch-clamp recordings of MDN, DNp17, and DopaMeander, neurons were visualized using w; 10xUAS-IVS-myr:GFP (Table S1). Experiments were conducted during daylight conditions and at room temperature. Flies were cold anesthetized at 4°C for immobilization, and the head and thorax were fixed to a pyramid-shaped fly holder. This leaves the legs and wings free, allowing walking and flight. The proboscis was fixed with glue to increase stability. Afterward, on the posterior side of the head, a small window was cut into the cuticle. Trachea, fat tissue, and the ocellar ganglion were removed to gain access to MDN, DopaMeander, and DNp17. Neurons of choice were visualized under a customized fluorescence microscope based on the SliceScope (Scientifica, Uckfield, UK). The neural sheath covering the location of the neurons was ruptured by local application of collagenase (0.025 % in extracellular saline) using a blunt patch-pipette. During preparation and experiments the fly brain was continuously perfused with carbonated extracellular saline (95% O2 and 5% CO2) containing: 103 mM NaCl, 3 mM KCl, 5 mM N-Tris (hydroxymethyl)methyl-2-aminoethane-sulfonic acid, 8 mM trehalose, 10 mM glucose, 26 mM NaHCO3, 1 mM NaH2PO4, 1.5 mM CaCl2 and 4 mM MgCl2 (adjusted to 273–275 mOsm, pH 7.3)^35^. Whole cell patch-clamp experiments were performed using patch-clamp electrodes (7-12 MΩ) filled with intracellular saline (40 mM potassium aspartate, 10 mM HEPES, 1 mM EGTA, 4 mM MgATP, 0.5 mM Na3 GTP, 1 mM KCl and 20 μM, adjusted to 260–275 mOsm, pH 7.3). Intracellular recordings were recorded in current clamp mode with a MultiClamp 700B amplifier (Molecular Devices, San Jose, CA, USA) and digitized (Digidata 1550B, Molecular Devices) with a 10 kHz low-pass filter and sampled at 20 kHz controlled by pCLAMP 11 Software Suite (Molecular Devices). Spikes were detected using MATLAB as described in^54^. In brief, thresholds were set manually in either the recording or the derivative of the recording to detect spikes. Accuracy of spike detection was confirmed for each experiment by manual inspection of three different sections of the recording (beginning, middle, and end).

### Tethered walking and flight

For experiments in which walking behavior was analyzed, a similar setup as described previously was used^54^. In brief, an air-supported ball (polypropylene, diameter: 6 mm, Spherotech GmbH, Fulda, Germany) was positioned under the tethered fly using a micromanipulator and a camera (acA1300-200um, Basler, Ahrensburg, Germany) perpendicular to the fly. The spherical treadmill was illuminated using an IR LED (850 nm), and the movement of the ball was captured by two camera sensors to reconstruct the fly’s walking trajectory. Walking was determined by changes in translational velocity and thresholds for the detection of onset and cessation of walking were set manually in MATLAB. Only trials in which the fly was resting before onset and after cessation of a walking bout were considered for analysis.

During flight experiments, the spherical treadmill was lowered, and flies either spontaneously initiated flight or it was induced by gentle air puffs. The termination of the flight was spontaneous. To analyze flight, we employed an adapted wingbeat tachometer that tracks wing movement through a light guide featuring a long pass filter (760 nm and above, model 969, University of Cologne (UoC), Animal Physiology Electronics Workshop, Cologne, Germany; based on a design from IO Rodeo https://github.com/iorodeo/light_sensor_boards). Flight trials were included for analysis if they featured at least 4 s of continuous flight in patch-clamp experiments (although they typically lasted much longer). In addition, trials were only included for further analysis if the inter-flight intervals were at least 10 s long. Flight onset and cessation were determined by extracting the timestamp of the first and last wingbeat in MATLAB for each flight bout.

### Antennal stimulation

For antennal mechanical stimulation, flies were cold anesthetized for immobilization and fixed to a pyramid-shaped fly holder as described above. In behavior-only experiments, flies were then ready to use, and further steps were conducted for experiments with simultaneous MDN recording, as described above. The mechanical stimulation was delivered by a precision high-speed linear actuator (Poker; Model 920, UoC, Animal Physiology Electronics Workshop). This device can generate a controlled linear and precision movement. A human hair was attached to the tip of the poker to have a very small area of effect. Before the experiment, the poker was placed approximately 2-3 mm away from the front of the head of the tethered fly. After manually confirming that the poker can deflect the antenna, the protocol was carried out consisting of 10 pokes, each lasting 2 seconds, with a 10-second interstimulus interval. The position of the poker and antennal deflection was visually inspected using two cameras (acA1300-200um, Basler, Ahrensburg, Germany), one frontal slightly from below and one perpendicular to the fly. For each fly, each antenna was only subjected once to the protocol, which was started alternately with the right or left antenna. The protocol was controlled by the pCLAMP 11 Software Suite (Molecular Devices) to ensure an exact alignment of poking events with behavioral data and neuronal activity.

### Free-walking assay

For optogenetic activation experiments, virgin female flies carrying 20x-UAS-CsChrimson-mvenus^88^ were mated with males from the SS02553-split-GAL4 line and Empty-split-GAL4 line. Post-eclosion, the offspring were reared on standard food coated with 300 µM all-trans-retinal for three to six days before the experiment. The walking activity of up to 20 flies was recorded using the behavioral setup named ‘Universal Fly Observatory’ (UFO), as described previously^37,89^. The UFO features a walking arena consisting of an inverted petri dish with a diameter of 100 mm as its base. To prevent escape, this base was covered with a watch glass of diameter 120 mm. The inside of the watch glass was coated with SigmaCote siliconizing reagent (cat. no. SL2; Sigma-Aldrich, RRID: SCR_008988) to prevent flies from walking upside down on the watch glass. The arena was illuminated by a ring of 60 infrared (IR) LEDs (870 nm) arranged concentrically around the arena. A diffuser was placed along the inner side of the IR LED ring to ensure uniform illumination across the entire arena. The arena was filmed from below via a surface mirror positioned at a 45° angle below the arena and a camera (Basler acA1300-200um, Basler AG, Ahrensburg, Germany) with a spatial resolution of 0.08 mm/pixel and a temporal resolution of 20 Hz. The camera’s lens was equipped with an IR-pass filter (lower cut-off frequency: 760nm) that only permits the IR light to pass through to ensure a strong contrast between the background (black) and the flies (bright). A second ring of 60 red LEDs (625 nm) positioned above the IR LED ring provided the light necessary to activate CsChrimson for optogenetic activation transiently. The entire UFO setup was enclosed within a box, together with a second UFO, which included two visible white light sources placed above the arenas for controlled light conditions.

Before each experiment, approximately twenty flies were transferred to an empty vial and anesthetized on ice for two minutes. Subsequently, flies were transferred to the UFO arena. They were given an acclimatization period of ten minutes to explore the arena before starting the experiment. We activated the SS02553-split-GAL4 line using five different intensities of red light(0.41, 1.35, 2.28, 3.19 and 4.07 mW/cm^2^). For each intensity, nine activation cycles were carried out. One activation cycle consisted of 10s of stimulation followed by a 60s interstimulus interval. A five minute window was provided between different intensities. All experiments were conducted under a background of white light at an intensity of 0.11mW/cm^2^. For each experiment, experimental and control flies were recorded simultaneously in two adjacent but independently controlled UFOs in the same enclosure. Both the protocol and video acquisition were controlled by custom-written MATLAB code.

### Tracking and analysis of walking trajectories

Movement of flies was tracked using the TRex software^83^. The tracking data were extracted from the videos on a frame-by-frame basis for further analysis in MATLAB. All tracking data was manually reviewed and identity switches were corrected manually.

Extracted and corrected fly coordinates (center of mass) and speed was further analysed in MATLAB. We only examined data around the 10 s optogenetic stimulations (2 seconds before stimulus onset until 2 s after stimulus offset).

To analyze the mean speed, a moving average filter with a 0.5 s window was applied for data smoothing. For each fly, the median speed prior to stimulation (2 s) was subtracted. The mean speed during activation was calculated in a time window of 8 s starting at the onset of the light pulse. The light off period at the end of the optogenetic stimulus was ignored because the sudden decrease in light occasionally leads to strong activity increases in flies, independent of genotype. Then, we computed the mean speed across trials for each fly to obtain the average speed distribution of the population.

To analyze turning behavior we simplified each fly path by collapsing trajectory time-points closer than 0.5 mm into a single point spanning the same time window. Then, we computed the complementary angle for each triplet of points that compose the trajectory to obtain a 0° angle when the fly walks straight and a ±180° in case of a turn in the opposite direction. We computed the cumulative angle for each fly summing the absolute values of the angles over time. The mean cumulative angle for each fly was computed in a time frame of 8 s from the beginning of the activation.

We calculated the total distance traveled by the fly and the distance between the initial and final points of the trajectory during the time window 0-8 s. Then, we used the mean across 9 trials of these two variables as a metric to analyze path tortuosity. A higher ratio between total distance and start-to-end distance is indicative of frequent directional changes, and a ratio near 1 suggests straight walking.

### Optomotor stimulation assay for silencing experiments

The UFO setup was placed in the center of a modular LED display adapted from Reiser et al. forming a cylindrical arena composed of 100 8×8 Blue (Φ3.0mm, 470nm) LED panels (KWM-30881ABB/ KWM-30881CBB, Luckylight), arranged in a 5×20 grid^90^. Each square panel has 32.0 mm height, forming a cylindrical display of 16 cm height and approximately 20 cm diameter around the UFO setup. To control the LED displays, we used the MATLAB script PControl (https://reiserlab.github.io/Modular-LED-Display/)^90^. Walking activity was promoted by presenting a pattern consisting of alternating 5 LED wide (15mm on the display) vertical bars the full height of the display. Intensity of the bars was set on level 2 of the PControl options (0 to 7 levels). The day before the experiment, 3-4 day old flies were starved overnight to increase their drive to walk^37^. On the test day, flies were briefly cold anesthetized and 4 of them were transferred to the arena. Males and females were tested separately from each other. Each experiment consisted of two phases of 4 min each. First, an adaptation phase where bars were presented as a stationary pattern, followed by a second phase in which the pattern rotated counterclockwise at a speed of 50 LEDs displacement per second (15 cm/s on the screen).

Flies were tracked and data loaded in MATLAB as previously described. For each video, we examined 10 time windows of 10 seconds, each one separated by a 10 seconds interval. Data were smoothed and analyzed as before but were not baseline subtracted. Mean speed, cumulative angle, total distance and start-to-end distance were analyzed over the full duration of each 10 seconds window and then averaged across the 10 time windows.

### Stochastic targeting of cells using SPARC

To determine which neuron types labelled by the SS02553-split-GAL4 line drive the walking phenotype, we used the genetic tool Sparse Predictive Activity through Recombinase Competition (SPARC) to achieve stochastic expression of CsChrimson in these neuron types in a controllable manner. We used two different variants of SPARC2-CsChrimson (i-intermediate,∼15% and d-dense,∼50%) to increase the diversity of stochastic expression patterns across flies. By expressing nSyb-PhiC31 recombinase pan-neuronally along with one of the two SPARC2-CsChrimson::tdTomato variants and SS02553-split-GAL4 we generated flies that expressed CsChrimson::tdTomato in different fractions of the 8 neuron types labelled by the SS02553-split-GAL4 line. Individual female flies were then optogenetically activated in the UFO using the protocol mentioned above using red light at an intensity of 4.07 mW/cm^2^. Flies that exhibited any walking phenotype were subjected to dissection and immunostaining to determine which neuron types were expressed. Randomly selected flies from the same genetic crosses were chosen as potential control flies, and also dissected and immunostained. Walking behaviour was tracked and analysed as described above for SS02553>CsChrimson.

### Immunohistochemistry and image acquisition

For immunolabeling of SS02553>SPARC, flies were anesthetized on ice and then transferred to 1.8mL 4% paraformaldehyde in 0.1 M phosphate buffered saline containing 0.5% of Triton-X100 (PBT). The flies were fixed in this solution for 6 h on a shaker at RT followed by three 15 min wash steps in PBT. Individual fly brains and VNCs were then dissected in a SYLGARD dish after immersing the fly briefly in 70% ethanol. Dissected brains were then washed three times in PBT for 15 min followed by blocking in 10% normal goat serum in 0.1M PBT (blocking buffer), overnight at 4°C. Samples were then incubated for 2 d at 4°C in primary antibody solution in blocking buffer. For staining tdTomato, we used anti-RFP diluted to 1:1000 and anti-nc82 diluted to 1:500 to label neuropils as background. Afterwards, samples were washed in PBT three times for 15 min each, followed by overnight incubation in secondary antibody solution at 4°C containing AlexaFluor 555 goat anti rabbit IgG (H+L) diluted to 1:200 and AlexaFluor 635 goat anti-mouse IgG diluted to 1:400. Finally, samples were washed three times in PBT, transferred to PBS and then mounted in Vectashield Antifade Mounting Medium.

For intracellular stainings with Neurobiotin, we used 0.5% Neurobiotin tracer (VEC-SP-1120-20, Vector Laboratories Inc., CA, United States) dissolved in intracellular saline. To fill DopaMeander, we injected current for about 10 min at the end of an experiment (20 pA at 120 Hz). Flies were then immediately transferred into 4% paraformaldehyde in 0.1 M phosphate buffered saline and fixed for 3 h on a rotorad at room temperature. Subsequently, brains were dissected, preincubated and stained using primary antibodies (anti-GFP, 1:200; anti-nc82, 1:50) as described above. Samples were washed three times in PBT for 15 min and incubated in secondary antibodies (AlexaFluor 635 goat anti-mouse, 1:400; AlexaFluor 488 goat anti-chicken, 1:200) and 1:500 Streptavidin Alexa Fluor 568 conjugate for three days at 4°C. Samples were then washed and embedded as described above.

Image acquisition was performed with a confocal laser-scanning microscope, Leica TCS SP8 WLL through the Leica Application Suite (LASX) using an HC PL APO 20x/0.75 IMM objective. Fluorescence was detected using suitable lasers with a resolution of 1024×1024 pixels, in serial stacks. Images were scanned sequentially to reduce dye cross-excitation. Final image processing was done in Fiji.

### Connectivity analysis

All analyses were performed using the FlyWire connectomics dataset (FAFB v783^40^) using synaptic predictions from Buhmann et al. 2021^91^. Connectivity between DopaMeander neurons (Figure S3B) was visualized using the shortest path length tool in FlyWire Codex, with a synapse threshold of 5 (https://codex.flywire.ai/app/path_length?dataset=fafb). All other plots were generated using custom R scripts interacting with the *natverse* and *fafbseg* packages^92^. Synaptic input and output profiles of DopaMeander were grouped either by their predicted neurotransmitter^42^, cell type, or superclass^11^. Synaptic inputs to AN_AVLP_51 neurons were pooled together and grouped by cell type (Figure S3E). To generate the connectivity graph in Figure 4D, an all-by-all connectivity matrix was retrieved for BPNs^14^, DNp17, MDNs, and descending neurons connected to DopaMeander. Neurons in this matrix were grouped by cell type. The matrix was imported into Gephi (v0.10.1) for graph visualization. Autapses and weak connections (<5 synapses) were omitted for clarity.

### Linear-Nonlinear models

For modelling of free locomotion features based on the neuronal activity of MDN, DopaMeander and DNp17, we implemented linear-nonlinear (LN) models. Neural activity and locomotion features were downsampled from 20 kHz to 200 Hz. All models were trained with 65% of the data, and the remaining 35% were held out for quantifying model performance. The LN models consisted of a temporal filter with a duration of 3s, estimated using time-delay embedding of the neural activity with a basis of 20 raised cosine functions to reduce the number of parameters (from 600 time bins to 20 basis functions) and to enforce smooth filters. We then fitted a linear model with L1 regularization to the corresponding locomotion feature to the time-delay-embedded inputs, and selected the optimal regularization strength by cross-validation (*LassoCV* from *scikit-learn*^87^). The nonlinear transformation was estimated from the training data predictions (using *binned_statistics* from the *scipy* python package^86^), and was applied to the filtered neural activity to generate the final model predictions. The LN model performance is the *Pearson R* score between the predictions and the test data.

Locomotion features were decomposed into unprocessed (raw), positive (pos), negative (neg) and absolute (abs) values. This was done by clipping the corresponding velocity at zero, or applying an absolute value function. For angular velocity, negative represents clockwise turning, and for translational velocity, negative represents backward walking.

For the rolling correlation analysis (Figure S5C-H) we calculated the model performance and locomotor features in time windows of 2 s.

For all pairwise comparisons of model performances (Figure 6D, F, H, J), we assessed the normality of each group using the Shapiro-Wilk test. If both groups were normally distributed, we employed independent t-tests for unequal variances. In cases where normality was not observed, we used a two-sided Mann-Whitney U test. All p-values are presented following Bonferroni correction for multiple comparisons.

To control for correlations between locomotor features when analyzing LN model performance for specific locomotor features, we “regressed out” particular aspects of locomotion^93^, Fig. S5B). We first removed shared variance between absolute translational and angular velocities, by regressing one against the other (absolute translational against angular velocity, and vice versa). The residuals of the regression capture the unique variance unexplained by the other variable and were then used to train individual LN models, following the same procedure as above.

### Quantification and statistical analysis

Statistical analysis was performed in MATLAB and python. The test used and relevant statistical parameters are reported in the respective figures, captions, Tables S2-4, or the main text. If not otherwise noted, values are presented as medians and interquartile ranges (IQR) throughout the text and figure legends.

## Acknowledgments

We thank our dear friend and colleague, the late Konrad Öchsner (JMU) for technical support, Michael Dübbert (UoC) and Thomas Walter (JMU) for help developing setups and electronics, Karen Hibbard (Janelia) and Salil Bidaye (MPFI) for sharing reagents, Charlotte Förster for sharing infrastructure, members of the Ache Lab and Till Bockemühl (UoC) for stimulating discussions, and Chris J. Dallmann for feedback on the manuscript. This work was supported by grants from the Deutsche Forschungsgemeinschaft (DFG, German Research Foundation) to J.M.A. via the Emmy Noether program (DFG AC 371/1-1) and the Next Generation Networks for Neuroscience (NeuroNex) research grant C^3^NS (DFG AC 371/2-1). J.C. is supported via the Emmy Noether program (DFG CL 596/1-1) and the ERC (Starting Grant NeuSoSen 851210); A.B. is supported by NeuroNex grant C^3^NS (DFG BU 857/15), DFG CRC 1451 (SFB1451/1 and 431549029), and is a member of the Nordrhein-Westfalen (NRW) network iBehave; M.E. is supported by a DFG Walter-Benjamin Fellowship (DFG ER 1168/1-1).

## Author contributions

SD performed MDN recordings, MDN-related behavioral experiments, and analyzed electrophysiology datasets; SL performed DopaMeander recordings, DNp17 recordings, and related analyses; FMI performed UFO and DopaMeander SPARC experiments and behavioral tracking; APM developed LN models; FCM analysed all UFO data; ME performed connectome analysis, AMD recorded MDN visual responses, EAG performed DopaMeander silencing experiment, AB facilitated DopaMeander silencing experiments and development of antennal stimulator, JC developed LN models, JMA conceived of the project, designed experiments, acquired funding, and supervised the overall project.

Conceptualization: JMA

Data Curation: SD, SL, FMI, EAG, FCM

Formal Analysis: SD, SL, APM, FCM, ME

Funding Acquisition: ME, JC, AB, JMA

Investigation: SD, SL, FMI, AMD, ME, APM, EAG

Methodology: SD, SL, JC, JMA

Project Administration: JMA

Resources: AB, JMA

Software: SD, SL, APM, FCM

Supervision: AB, JC, JMA

Validation: AMD, FCM

Visualization: SD, SL, APM, FCM, ME, JMA

Writing – Original Draft Preparation: SL, SD, and JMA with help from APM and ME.

Writing – Review & Editing: SL, SD, FMI, APM, FCM, and JMA with help from all authors.

## Inclusion and diversity

One or more of the authors of this paper self-identifies as an underrepresented ethnic minority in their field of research or within their geographical location. One or more of the authors of this paper self-identifies as a gender minority in their field of research. We support inclusive, diverse, and equitable conduct of research.

## Supplemental information

**Figure S1:**
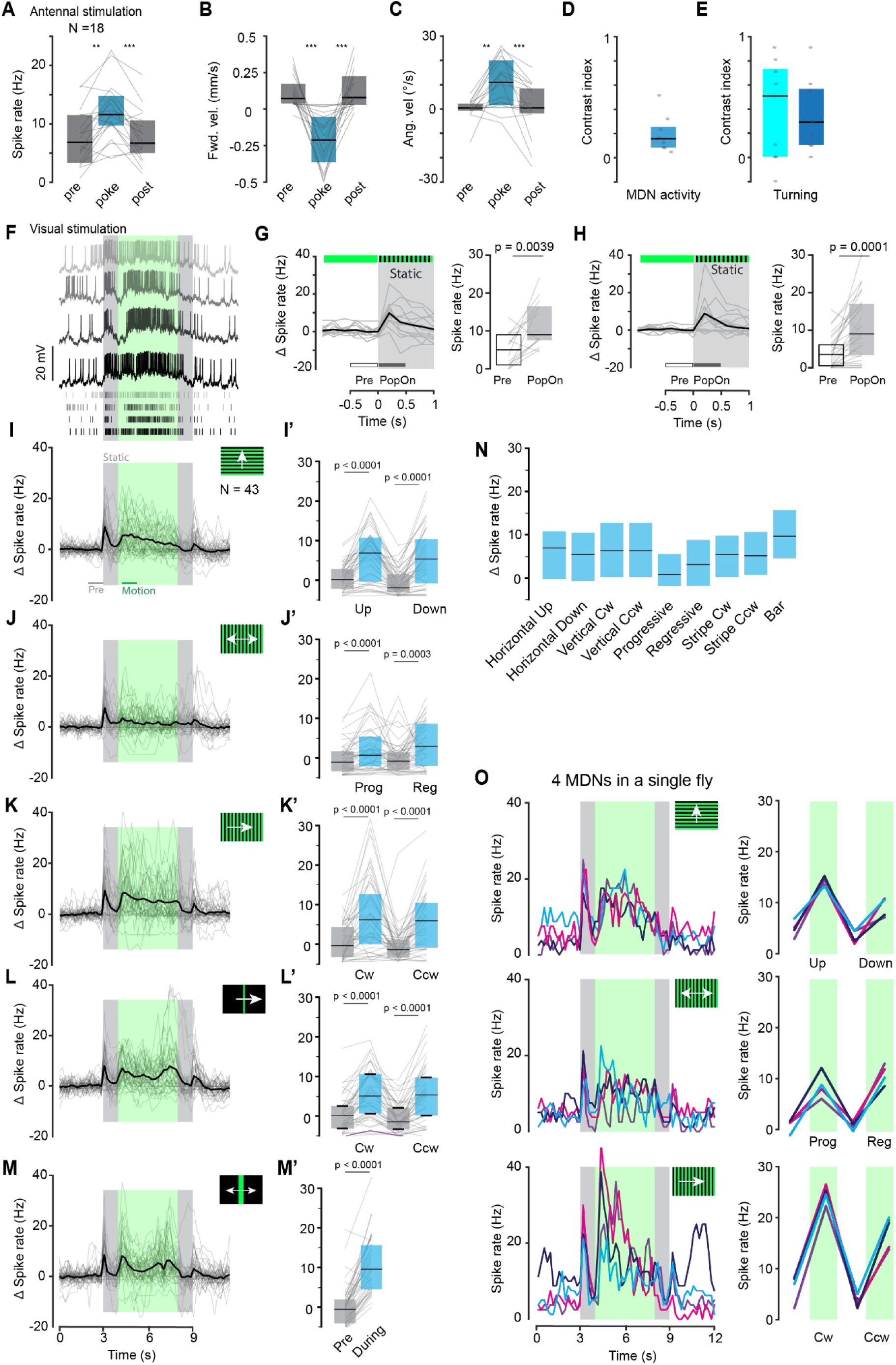
Visual and mechanosensory responses of MDN. A) Quantification of MDN spike rate during antennal touch (pooled across left and right antenna). Spike rate was averaged in a 500 ms window before (pre), during the first 500 ms of antennal touch (poke), and the 500 ms after antennal touch (post). Data shown as median ± IQR. B) Quantification of forward velocity during MDN recordings in response to antennal touch. Forward velocity was average during 1 s before (pre), during the first 1s of the stimulus (poke), and 1s after the stimulus. Other details as in (A). C) Angular velocity changes during MDN recording in response to antennal touch. Other details as in (B). D) Selectivity of MDN response for a preferred antennal side. Calculated using contrast analysis comparing spike rates from stimulation of the preferred antenna (stronger response) vs. the unpreferred antenna (weaker response) for each MDN recording (see Figure 1, grey dots, N=9). Contrast was calculated as (A-B)/A+B), so that it would yield 1 or −1 for an MDN that responded exclusively to one antenna. E) Contrast analysis of turning behavior in response to antennal touch. Calculated using contrast analysis comparing left and right turns in response to left (light blue) and right (dark blue) antennal touch for each fly (grey dots). Note, that the average behavioral response sensitivity is much higher than the potential maximal selectivity of the MDN population calculated in D. F) Example recording of MDN membrane potential (Vm) and spike events during visual stimulation (upward motion as in I). Four presentations of the same stimulus shown for one MDN. G) Change in MDN spike rate in response to a static vertical grating (PopOn) in intact flies (N = 14 MDNs). Boxplots show data as median ± IQR with lines representing individual flies. P-values were calculated using the Wilcoxon signed-rank test. H) Same as in (G) for flies with ablated antennae (N = 29 MDNs). I) Change in MDN spike rate in response to a horizontal grating pattern moving up. I’) Quantification of change in MDN spike rate in response to horizontal grating pattern moving up or down. Boxplot shows data as median ± IQR with lines representing individual flies. P-values were calculated using the Wilcoxon signed-rank test. J) to M) Sames as in I) but for progressive and regressive visual motion (J and J’), vertical grating rotating clockwise (cw) or counter clockwise (ccw) (K and K’), a single stripe moving cw or ccw (L and L’) and an expanding bar presented in front of the animal (M). N) Comparison of change in spike rate in response to widefield visual motion and moving object stimuli. O) Change in spike rates of 4 MDNs recorded subsequently in a single fly responding to a horizontal grating moving up or down, progressively and regressively, and a vertical grating moving cw or ccw. Note that all visual responses in F-O were recorded in tethered, non-walking flies with glued legs to analyze visual responses independent of behavioral responses.

**Figure S2:**
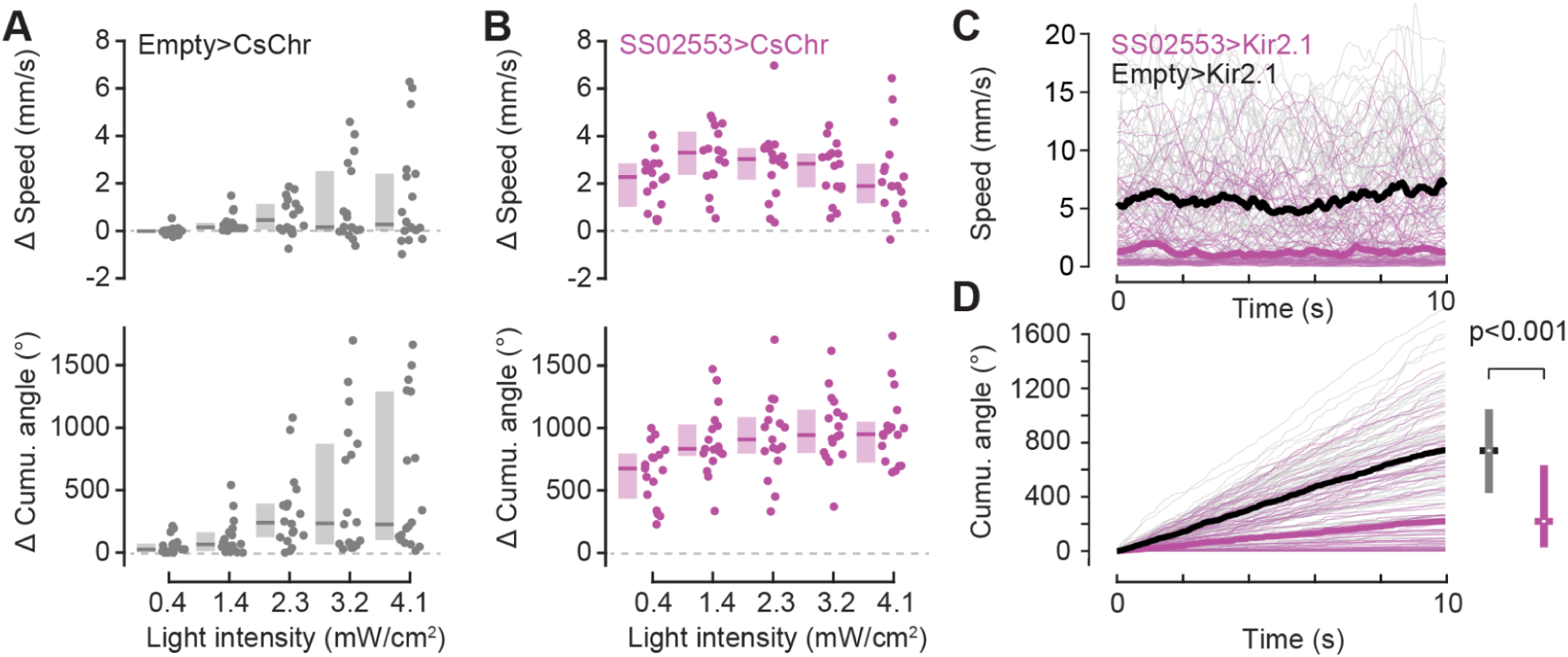
Activation of SS02553 increases walking speed and curve walking whereas silencing SS02553 decreases curve walking. A) Optogenetic stimulation protocol had minor effects on walking speed (top) and curve walking (bottom) in empty control flies. N=18 flies, 9 trials per intensity. B) Optogenetic activation of SS02553 increased walking speed (top) and curve walking (bottom) across activation intensities. Increasing the activation intensity had little effect on the speed or cumulative angle. N=17 flies, 9 trials per intensity. C) Silencing SS02553 strongly reduces forward walking speed in starved flies exposed to an optomotor stimulus. empty>Kir2.1 N=80, SS02553>Kir2.1 N=80. See main Figure 3 for statistics. D) Silencing SS02553 significantly reduces curve walking, i.e., the cumulative angle over time, compared to empty>Kir2.1 controls. P-value calculated via Wilcoxon rank sum test. In all panels, boxplots show median and IQR.

**Figure S3:**
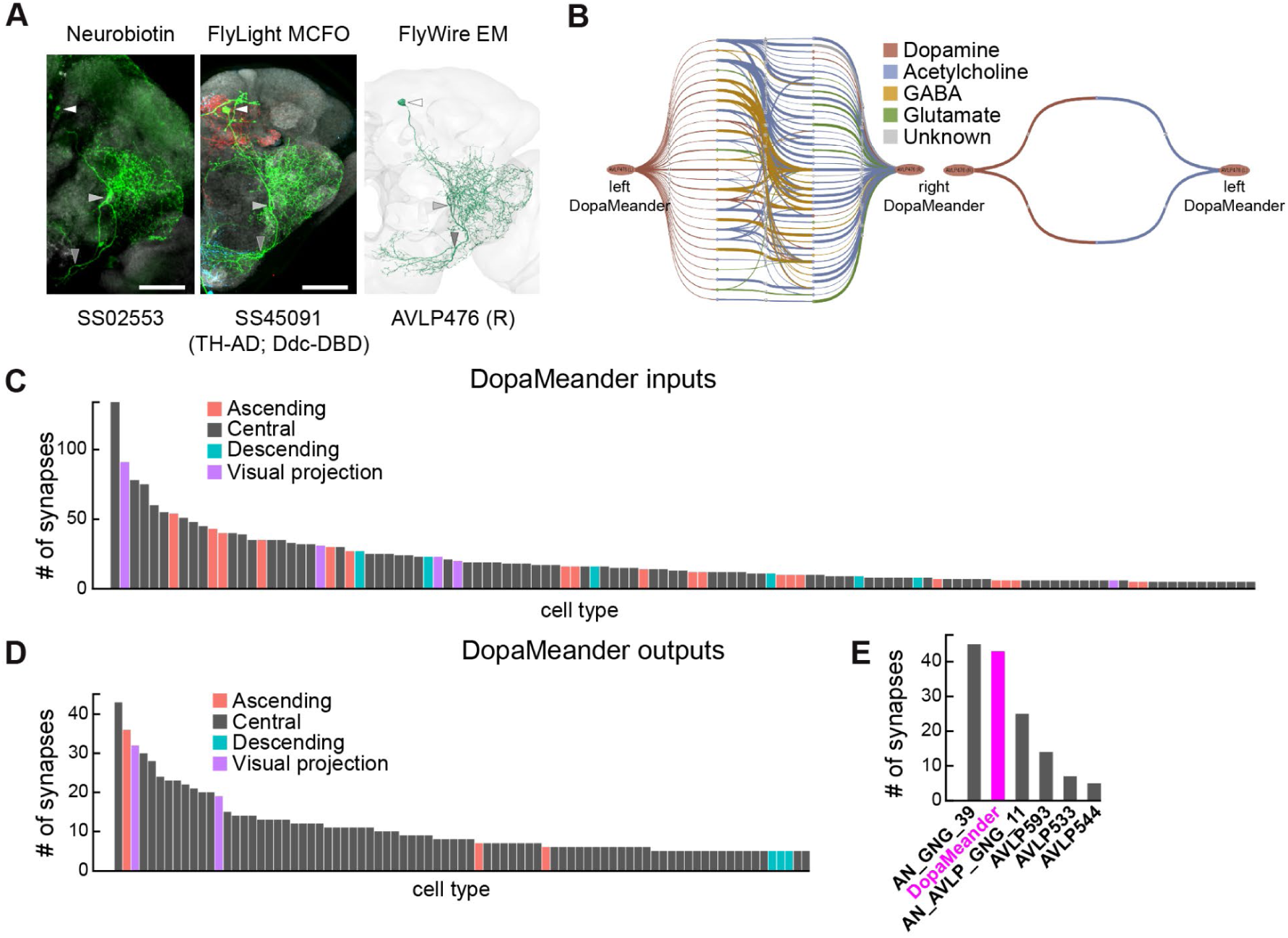
Synaptic connectivity profile of DopaMeander. A) DopaMeander anatomy revealed by different approaches. Left, neurobiotin injection into DopaMeander cell in SS02553 driver line. Middle, MCFO image of dopamine-specific SS45091 driver line. Right, FlyWire electron microscopy (EM) reconstruction of DopaMeander neuron. White, light gray and gray arrowheads indicate the soma and the main neurites entering AVLP and GNG regions, respectively. Scale bar: 50 µm B) Left, shortest path of synaptic connectivity from left DopaMeander to right DopaMeander. The line thickness depicts synaptic strength and the color depicts the predicted neurotransmitter. Right, shortest path from right DopaMeander to left DopaMeander. C) Input profile of DopaMeander, grouped by cell type and colored by neuron superclass. D) Output profile of DopaMeander, grouped by cell type and colored by neuron superclass. E) Input profile of AN_AVLP_51. DopaMeander is highlighted in magenta.

**Figure S4:**
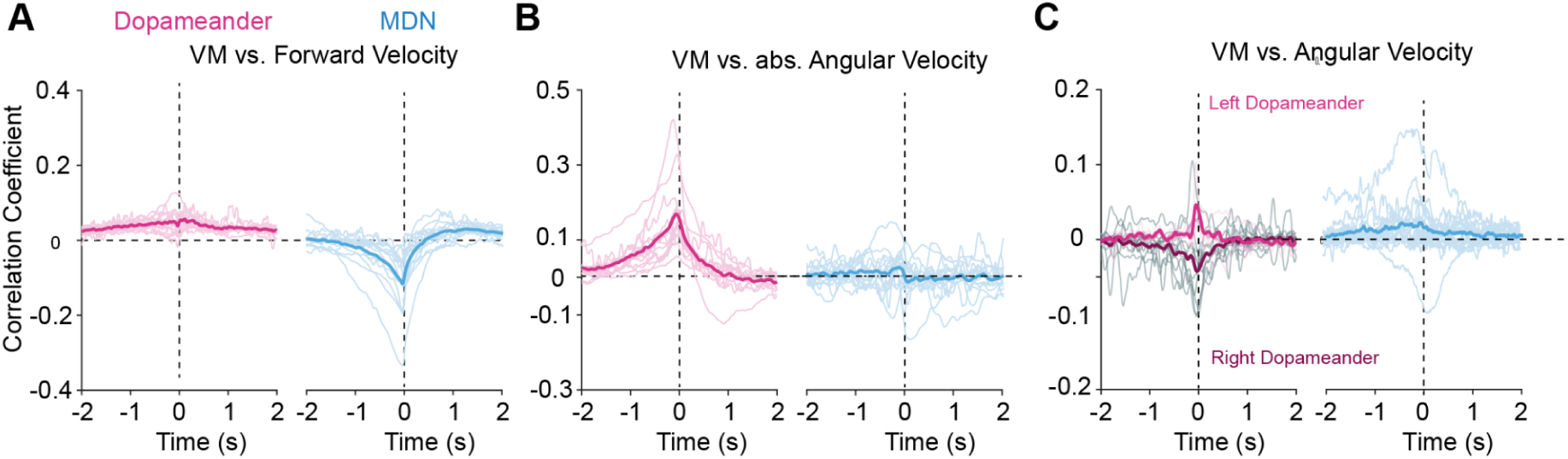
Correlation of DopaMeander and MDN membrane potential and walking parameters. A) Cross-correlation analysis for DopaMeander (magenta, N=16) and MDN (blue, N=15) membrane potential (VM) and backward velocity, both in 250 ms bins. All data sets were z-scored before correlation analysis. B) Same as (A) but for absolute angular velocity, i.e., turning. C) Same as (A) but for angular velocity, i.e., left or right turning. Left (light) and right DopaMeander (dark) populations could be distinguished during the recording

**Figure S5.**
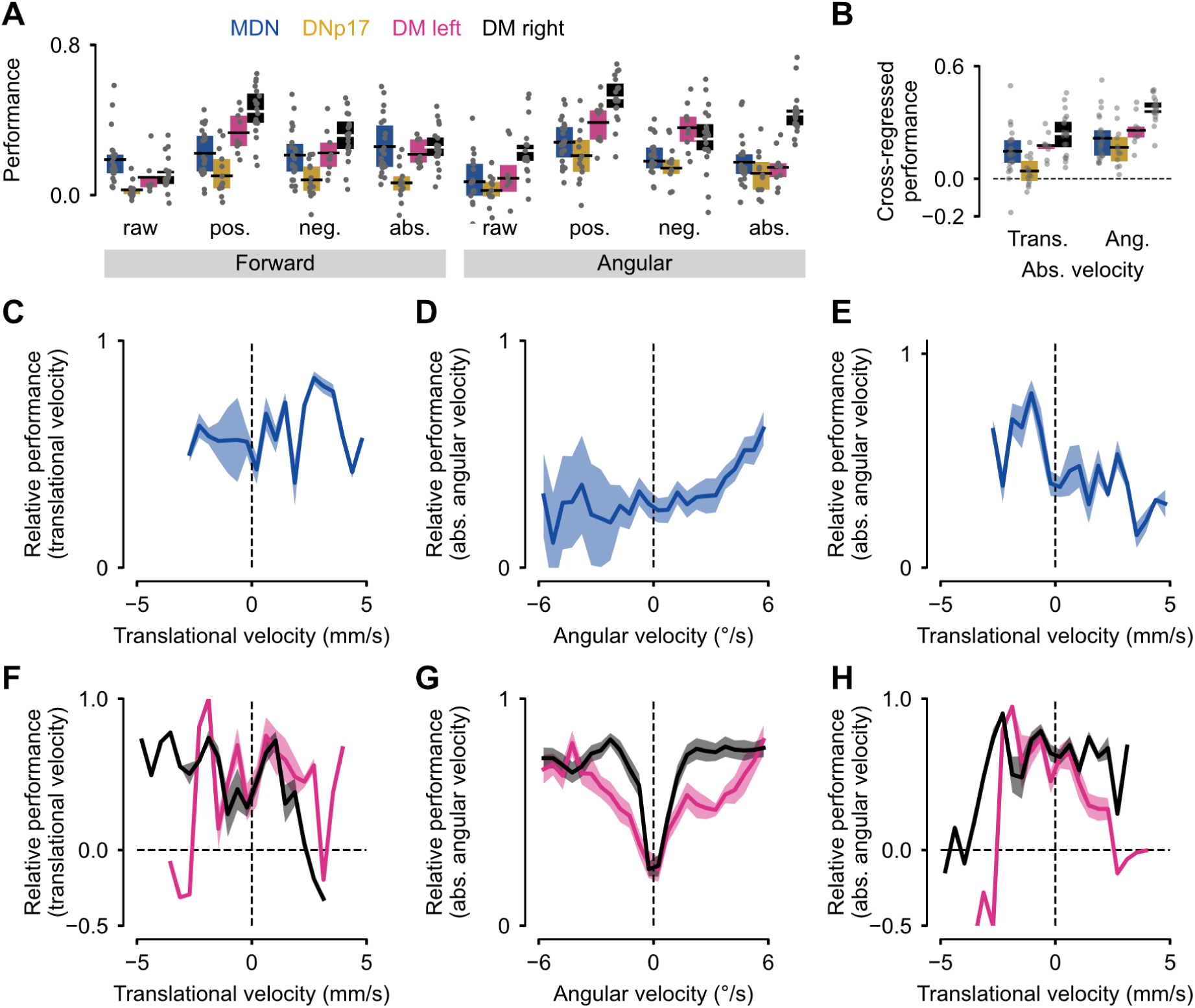
All LN model performances, and relevant rolling correlations. A) DNp17, DopaMeander and MDN tested across all locomotion components. B) Residualization (as in Garcia et al., 2020) across the absolute component of both translational and angular velocities. Models predict a significant unique component from the variance of translational and angular locomotion, and not just the correlated components from common locomotion across features. C-H) Normalized rolling correlation between MDN (C-E) and DopaMeander (F-H) prediction of translational velocity (C,F) and absolute angular velocity (D-E, G-H), against the rolling mean (2swindow) of translational velocity (C,E,F,H) and angular velocity (D,G). Solid line: mean; shaded area: standard error of the mean.

**Video S1. SS02553 walking phenotype elicited by optogenetic activation.**

Representative video showing the walking phenotype of SS02553>CsChrimson and parallel empty>CsChrimson control experiments in the UFO arena. Videos show 5s before and 5s after each trial of activation at red light intensity 4.1mW/cm^2^. White box on top left indicates light ON. Video is shown at 2x original speed.

**Table S1:** Fly genotypes used by figures and videos, related to Figures 1-7, Figures S1-S7 and Videos S1.

| Figure | Panel | Genotype |
| --- | --- | --- |
| Figure 1 | A-B, D-E, G-J | w[*]/-;VT044845-DBD/+; VT050660-AD/P(10XUAS-IVS-GFP)attp2 |
| Figure 1 | D-E | w+;VT044845-DBD/+;VT050660-AD/pJFRC49-10XUAS-IVS-eGFPKir2.1(attP2) |
| Figure 2 | B-H | P(y[+t7.7]w[+mC]=20XUAS-IVS-CsChrimson.mVenus)attP18/+; P(y[+t7.7]w[+mC]=p65.AD.Uw)attP40/+; P(y[+t7.7] w[+mC]=GAL4.DBD.Uw)attP2/+ |
| Figure 2 | B-H | P(y[+t7.7]w[+mC]=20XUAS-IVS-CsChrimson.mVenus)attP18/+;P(y[+t7.7]w[+mC]=R67E08-p65.AD)attP40/+; P(y[+t7.7] w[+mC]=VT025789-GAL4.DBD)attP2/+ |
| Figure 2 | I-L | w+; P(y[+t7.7] w[+mC]=p65.AD.Uw)attP40/+; P(y[+t7.7] w[+mC]=GAL4.DBD.Uw)attP2/pJFRC49-10XUAS-IVS-eGFPKir2.1 (attP2) |
| Figure 2 | I-L | w-; P(y[+t7.7] w[+mC]=R67E08-p65.AD)attP40; P(y[+t7.7] w[+mC]=VT025789-GAL4.DBD)attP2/pJFRC49-10XUAS-IVS-eGFPKir2.1 (attP2) |
| Figure 3 | A-B | w[*]/-; P(y[+t7.7] w[+mC]=R67E08-p65.AD)attP40; P(y[+t7.7] w[+mC]=VT025789-GAL4.DBD)attP2/P(10XUAS-IVS-GFP)attp2 |
| Figure 3 | C-E | P(y[+t7.7]w[+mC]=nSyb-IVS-PhiC31)attP18/+;P(y[+t7.7] w[+mC]=R67E08-p65.AD)attP40/TI(20XUAS-SPARC2-D-Syn21-CsChrimson::tdTomato-3.1))CR-P40;P(y[+t7.7] w[+mC]=VT025789-GAL4.DBD)attP2<br>P(y[+t7.7]w[+mC]=nSyb-IVS-PhiC31)attP18/+;P(y[+t7.7] w[+mC]=R67E08-p65.AD)attP40/TI(20XUAS-SPARC2-I-Syn21-CsChrimson::tdTomato-3.1))CR-P40;P(y[+t7.7] w[+mC]=VT025789-GAL4.DBD)attP2 |
| Figure 4 | - | - |
| Figure 5 | A, C-F | w-; P(y[+t7.7] w[+mC]=R67E08-p65.AD)attP40; P(y[+t7.7] w[+mC]=VT025789-GAL4.DBD)attP2/pJFRC49-10XUAS-IVS-eGFPKir2.1 (attP2) |
| Figure 5 | B, C-F | w[*]/-;VT044845-DBD/+; VT050660-AD/P(10XUAS-IVS-GFP)attp2 |
| Figure 6 | A-G | w[*]/-;VT044845-DBD/+; VT050660-AD/P(10XUAS-IVS-GFP)attp2 |
| Figure 6 | A-C, H-K | w[*]/-; P(y[+t7.7] w[+mC]=R67E08-p65.AD)attP40; P(y[+t7.7] w[+mC]=VT025789-GAL4.DBD)attP2/P(10XUAS-IVS-GFP)attp2 |
| Figure 7 | A-B | w[*]/-;VT044845-DBD/+; VT050660-AD/P(10XUAS-IVS-GFP)attp2 |
| Figure 7 | C-D | w[*]/-; P(y[+t7.7] w[+mC]=R67E08-p65.AD)attP40; P(y[+t7.7] w[+mC]=VT025789-GAL4.DBD)attP2/P(10XUAS-IVS-GFP)attp2 |
| Figure S1 | A-O | w[*]/-;VT044845-DBD/+; VT050660-AD/P(10XUAS-IVS-GFP)attp2 |
| Figure S2 | A | P(y[+t7.7]w[+mC]=20XUAS-IVS-CsChrimson.mVenus)attP18/+;P(y[+t7.7] w[+mC]=R67E08-p65.AD)attP40/+; P(y[+t7.7] w[+mC]=VT025789-GAL4.DBD)attP2/+ |
| Figure S2 | B | P(y[+t7.7]w[+mC]=20XUAS-IVS-CsChrimson.mVenus)attP18/+; P(y[+t7.7] w[+mC]=p65.AD.Uw)attP40/+; P(y[+t7.7] w[+mC]=GAL4.DBD.Uw)attP2/+ |
| Figure S2 | C-D | w+; P(y[+t7.7] w[+mC]=p65.AD.Uw)attP40/+; P(y[+t7.7] w[+mC]=GAL4.DBD.Uw)attP2/pJFRC49-10XUAS-IVS-eGFPKir2.1 (attP2) |
| Figure S2 | C-D | w-; P(y[+t7.7] w[+mC]=R67E08-p65.AD)attP40; P(y[+t7.7] w[+mC]=VT025789-GAL4.DBD)attP2/pJFRC49-10XUAS-IVS-eGFPKir2.1 (attP2) |
| Figure S3 | A | w[*]/-; P(y[+t7.7] w[+mC]=R67E08-p65.AD)attP40; P(y[+t7.7] w[+mC]=VT025789-GAL4.DBD)attP2/P(10XUAS-IVS-GFP)attp2 |
| Figure S4 | A-C | w[*]/-;VT044845-DBD/+; VT050660-AD/P(10XUAS-IVS-GFP)attp2 |
| Figure S4 | A-C | w[*]/-; P(y[+t7.7] w[+mC]=R67E08-p65.AD)attP40; P(y[+t7.7] w[+mC]=VT025789-GAL4.DBD)attP2/P(10XUAS-IVS-GFP)attp2 |
| Figure S5 | A-C | w[*]/-; P(y[+t7.7] w[+mC]=R67E08-p65.AD)attP40; P(y[+t7.7] w[+mC]=VT025789-GAL4.DBD)attP2/P(10XUAS-IVS-GFP)attp2 |
| Figure S5 | A-B- C-E | w[*]/-;VT044845-DBD/+; VT050660-AD/P(10XUAS-IVS-GFP)attp2 |
| Figure S5 | A-B, F-H | w[*]/-; P(y[+t7.7] w[+mC]=R67E08-p65.AD)attP40; P(y[+t7.7] w[+mC]=VT025789-GAL4.DBD)attP2/P(10XUAS-IVS-GFP)attp2 |

**Table S2:** Average variance explained by LN models, and corresponding standard deviation.

|  |  |  |  | Average variance explained | Standard deviation |
| --- | --- | --- | --- | --- | --- |
| Neuron | Side | Velocity | Component |  |  |
| DopaMeander | Left | Angular | Absolute | 0.39 | 0.12 |
|  |  |  | Right | 0.15 | 0.10 |
|  |  |  | Left | 0.36 | 0.12 |
|  |  |  | Unprocessed | 0.09 | 0.18 |
|  |  | Translational | Absolute | 0.33 | 0.11 |
|  |  |  | Backward | 0.22 | 0.07 |
|  |  |  | Forward | 0.22 | 0.12 |
|  |  |  | Unprocessed | 0.09 | 0.10 |
|  | Right | Angular | Absolute | 0.52 | 0.12 |
|  |  |  | Right | 0.43 | 0.11 |
|  |  |  | Left | 0.30 | 0.17 |
|  |  |  | Unprocessed | 0.24 | 0.15 |
|  |  | Translational | Absolute | 0.45 | 0.13 |
|  |  |  | Backward | 0.26 | 0.11 |
|  |  |  | Forward | 0.32 | 0.13 |
|  |  |  | Unprocessed | 0.11 | 0.09 |
| MDN | Undefined | Angular | Absolute | 0.28 | 0.12 |
|  |  |  | Right | 0.18 | 0.13 |
|  |  |  | Left | 0.18 | 0.16 |
|  |  |  | Unprocessed | 0.07 | 0.15 |
|  |  | Translational | Absolute | 0.22 | 0.12 |
|  |  |  | Backward | 0.26 | 0.15 |
|  |  |  | Forward | 0.21 | 0.14 |
|  |  |  | Unprocessed | 0.19 | 0.14 |

**Table S3:** Statistical tests and p-values before and after correction in Figure 6.

| Group 1 | Group 2 | Panel | Test | p | p (bonf corrected) |
| --- | --- | --- | --- | --- | --- |
| 0 | 2 | D | mannwhitneyu | 8.67E-02 | 2.60E-01 |
| 2 | 3 | D | ttest | 3.98E-01 | 1.00E+00 |
| 1 | 2 | D | ttest | 3.03E-01 | 9.08E-01 |
| 0 | 3 | F | ttest | 3.75E-05 | 1.13E-04 |
| 1 | 3 | F | mannwhitneyu | 2.28E-02 | 6.84E-02 |
| 2 | 3 | F | mannwhitneyu | 2.84E-03 | 8.53E-03 |
| 2 | 4 | H | mannwhitneyu | 8.73E-01 | 1.00E+00 |
| 3 | 5 | H | ttest | 1.06E-01 | 4.24E-01 |
| 0 | 6 | H | mannwhitneyu | 3.32E-03 | 1.33E-02 |
| 1 | 7 | H | ttest | 2.50E-10 | 1.00E-09 |
| 2 | 4 | J | ttest | 1.19E-03 | 4.74E-03 |
| 3 | 5 | J | mannwhitneyu | 3.64E-03 | 1.45E-02 |
| 0 | 6 | J | ttest | 1.97E-03 | 7.89E-03 |
| 1 | 7 | J | ttest | 7.07E-07 | 2.83E-06 |

**Table S4:** Statistical tests used in Figure 1-3, 4, and 7.

| Two-sided Wilcoxon rank-sum test |  |  |  |
| --- | --- | --- | --- |
| Figure | Variable | Context | P-value |
| Figure 1E | Forward Velocity | Ctrl. vs. Silenced | 0.003 |
|  | Angular Velocity |  | 0.1893 |
| Figure 2D | Baseline-subtracted speed | Empty vs SS02553>CsChrimson | 2.08E-17 |
| Figure 2F | 0s-subtracted cumulative angle | Empty vs SS02553>CsChrimson | 6.23E-17 |
| Figure 2H | Tortuosity index | Empty vs SS02553>CsChrimson | 8.66E-07 |
| Figure 2J | Speed | Empty vs SS02553>Kir2.1 | 1.88E-09 |
| Figure 2L | Tortuosity index | Empty vs SS02553>Kir2.1 | 7.37E-09 |
| Figure 3E | Baseline-subtracted speed | Dopameander&DNp17>SPARC vs DNp17 without DM>SPARC | 0.013986014 |
| Figure 3H | Tortuosity index | Dopameander&DNp17>SPARC vs None>SPARC | 0.001069519 |
| Figure S2A-B | Baseline-subtracted speed | intensity 1 | 8.05E-07 |
|  |  | intensity 3 | 1.33E-06 |
|  |  | Intensity 5 | 2.57E-05 |
|  |  | Intensity 7 | 0.004304694 |
|  |  | Intensity 9 | 0.057724701 |
| Figure S2A-B | 0s-subtracted cumulative angle | intensity 1 | 4.73E-07 |
|  |  | intensity 3 | 6.79E-07 |
|  |  | Intensity 5 | 4.58E-05 |
|  |  | Intensity 7 | 0.003877858 |
|  |  | Intensity 9 | 2.59E-02 |
| Figure S2D | Cumulative angle | Empty vs SS02553>Kir2.1 | 2.52E-09 |

| Two-sided Wilcoxon signed-rank test |  |  |  |
| --- | --- | --- | --- |
| Figure | Variable | Condition | P-Value |
| Figure 1H | MDN spike rate | Right Pre vs. right poke | 0.0195 |
|  |  | Left Pre vs. left poke | 0.0078 |
| Figure 1I | Forward Velocity | Right Pre vs. right poke | 0.0039 |
|  |  | Left Pre vs. left poke | 0.0039 |
| Figure 1J | Angular Velocity | Right Pre vs. right poke | 0.0273 |
|  |  | Left Pre vs. left poke | 0.0273 |
| Figure 7B | MDN spike rate | Pre vs. Flight 1 | 0.03125 |
|  |  | Flight 2 vs. Post | 0.03125 |
| Figure 7D | DopaMeander spike rate | Pre vs. Flight 1 | 0.0234 |
|  |  | Flight 2 vs. Post | 0.0078 |
|  | DopaMeander VM | Pre vs. Flight 1 | 0.0156 |
|  |  | Flight 2 vs. Post | 0.0078 |

